# Common molecular mechanisms underlie the transfer of alpha-synuclein, Tau and huntingtin and modulate spontaneous activity in neuronal cells

**DOI:** 10.1101/2021.07.18.452825

**Authors:** Inês Caldeira Brás, Mohammad Hossein Khani, Eftychia Vasili, Wiebke Möbius, Dietmar Riedel, Iwan Parfentev, Ellen Gerhardt, Christiane Fahlbusch, Henning Urlaub, Markus Zweckstetter, Tim Gollisch, Tiago Fleming Outeiro

## Abstract

The misfolding and accumulation of disease-related proteins are common hallmarks among several neurodegenerative diseases. Alpha-synuclein (aSyn), Tau and huntingtin (wild-type and mutant, 25QHtt and 103QHtt, respectively) were recently shown to be transferred from cell-to-cell through different cellular pathways, thereby contributing to disease progression and neurodegeneration. However, the relative contribution of each of these mechanisms towards the spreading of these different proteins and the overall effect on neuronal function is still unclear.

To address this, we exploited different cell-based systems to conduct a systematic comparison of the mechanisms of release of aSyn, Tau and Htt, and evaluated the effects of each protein upon internalization in microglial, astrocytic, and neuronal cells. In the models used, we demonstrate that 25QHtt, aSyn and Tau are released to the extracellular space at higher levels than 103QHtt, and their release can be further augmented with the co-expression of USP19. Furthermore, cortical neurons treated with recombinant monomeric 43QHtt exhibited alterations in neuronal activity that correlated with the toxicity of the polyglutamine expansion. Tau internalization resulted in an increase in neuronal activity, in contrast to slight effects observed with aSyn. Interestingly, all these disease-associated proteins were present at higher levels in ectosomes than in exosomes. The internalization of both types of extracellular vesicles (EVs) by microglial or astrocytic cells elicited the production of pro-inflammatory cytokines and promoted an increase in autophagy markers. Additionally, the uptake of the EVs modulated neuronal activity in cortical neurons.

Overall, our systematic study demonstrates the release of neurodegenerative disease-associated proteins through similar cellular pathways. Furthermore, it emphasizes that protein release, both in a free form or in EVs, might contribute to a variety of detrimental effects in receiving cells and to progression of pathology, suggesting they may be exploited as valid targets for therapeutic intervention in different neurodegenerative diseases.

**Graphical abstract:** 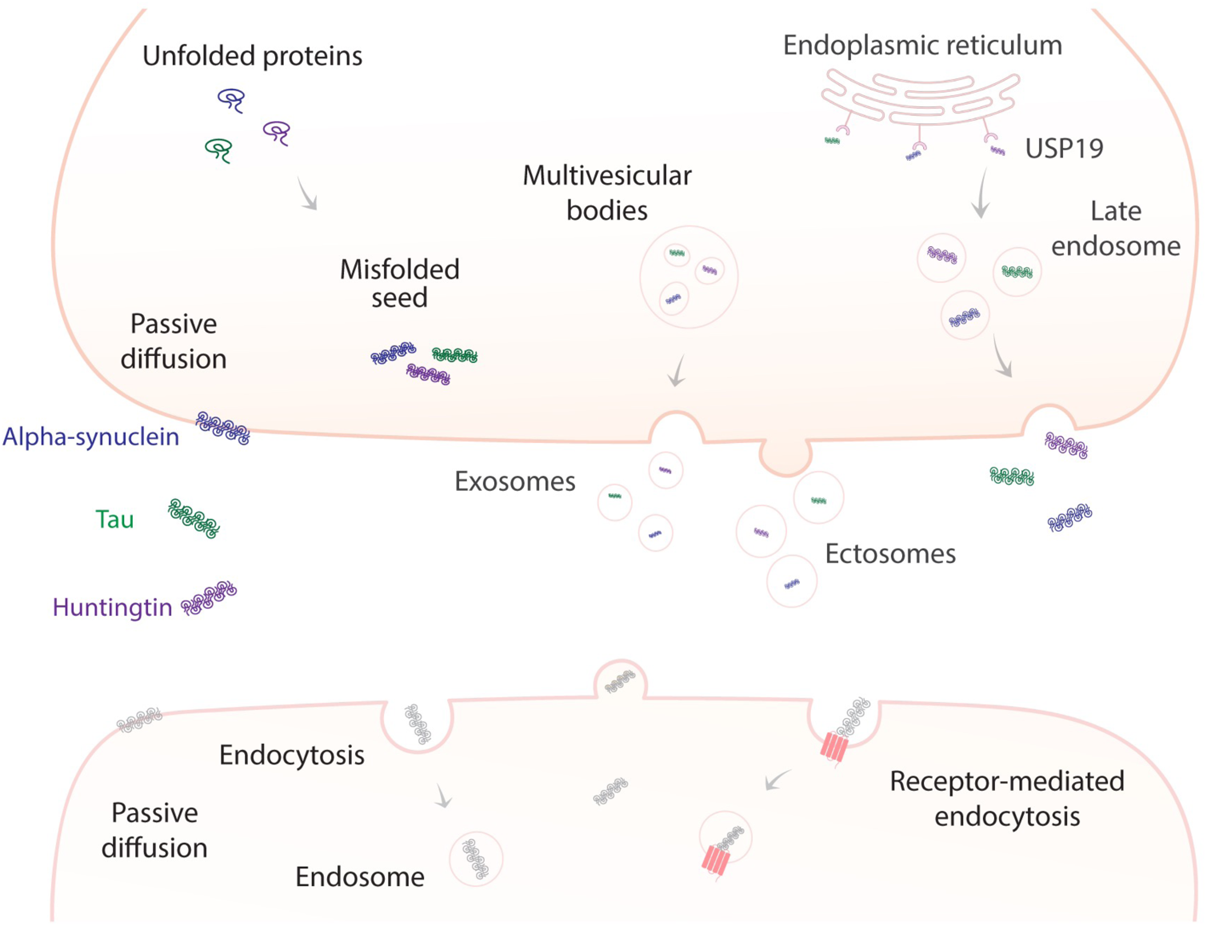

## Introduction

Neurodegenerative diseases are associated with the progressive loss of a variety of brain functions due to the loss of different types of neuronal cells. In spite of characteristic clinical manifestations, these disorders share common neuropathological features and cellular alterations, such as the aggregation and accumulation of disease-related proteins in relatively specific regions of the brain (Gan et al., 2018). Alpha-synuclein (aSyn)-containing aggregates are typical in Parkinson’s disease (PD) and other synucleinopathies, hyperphosphorylated Tau-containing inclusions are typical in tauopathies, and mutant huntingtin (Htt)-containing inclusions are typical in Huntington’s disease (HD) (Bras et al., 2020, Gotz et al., 2019, Tabrizi et al., 2020).

The progressive accumulation of protein pathology in different brain regions (Braak and Braak, 1991, Braak and Braak, 1995, Braak and Del Tredici, 2016, Braak et al., 2002, Braak et al., 2003) and the observation of aSyn Lewy body (LB) pathology fetal dopaminergic neurons grafted in the brains of PD patients led to the hypothesis that the progression of different neurodegenerative diseases may be correlated with the transfer of pathological proteins from sick cells to healthy cells (Li et al., 2008, Kordower et al., 2008). Consistently, injection of Tau aggregates into the brains of transgenic animals induces pathology along connected brain networks (Clavaguera et al., 2009, Iba et al., 2013, de Calignon et al., 2012, Liu et al., 2012). Furthermore, a similar process was hypothesized to occur also in monogenic forms of neurodegenerative diseases after the observation of mutant Htt aggregates within fetal striatal allografts in the brains of HD patients (Cicchetti et al., 2014).

The old brain is characterized by multi-morbidity, with the simultaneous accumulation of different types of protein pathology (Spires-Jones et al., 2017, Attems, 2017, Outeiro, 2021). In addition, abundant Tau-related pathology can be observed in the brains of PD and HD patients, suggesting that multiple proteins may jointly contribute to the pathophysiology of different neurodegenerative disorders (Cisbani et al., 2017, Fernandez-Nogales et al., 2014, Ornelas et al., 2020, Arima et al., 2000). Furthermore, the propagation of the pathological proteins between cells and across anatomical connected regions is consistent with the progression patterns described in different neurodegenerative diseases (Goedert et al., 2017, Bras and Outeiro, 2021, Alpaugh and Cicchetti, 2021). However, the precise molecular mechanisms underlying the spreading of pathology, and the relative contributions of each of them towards spreading, are still unclear. At a fundamental level, it is also unclear whether cells utilize similar pathways for releasing proteins, such as aSyn, Tau or Htt, and how the released proteins affect neighboring cells.

Several conventional and unconventional pathways have been implicated in the cell-to-cell transfer of proteopathic seeds (Peng et al., 2020, Bras and Outeiro, 2021, Alpaugh and Cicchetti, 2021). The conventional secretory pathway requires the presence of a signal peptide sequence in the secreted protein that is then translocated to the endoplasmic-reticulum (ER), and sorted through Golgi-derived vesicles that, ultimately, fuse with the plasma membrane, thereby releasing their content/cargoes to the extracellular space (Andrews, 2000, Ponpuak et al., 2015, Zhang and Schekman, 2013, Lee et al., 2004). Alternatively, cargoes can be sorted and released through unconventional pathways, resulting in the release of proteins in free forms, in extracellular vesicles (EVs) (Hill, 2019), or via tunnelling nanotubes, structures that enable the direct transfer of cargoes between connected cells (Abounit et al., 2016).

EVs have been shown to play various roles in the central nervous system (CNS), such as in intercellular communication, or the removal of toxic materials from the cell. In this context, they may also contribute to the transfer of pathogenic proteins in neurodegenerative diseases. EVs may be classified as exosomes or microvesicles (also known as ectosomes). They differ significantly in size, mechanism of biogenesis, and in protein, lipid, and nucleic acid content (Kalra et al., 2016, van Niel et al., 2018). Exosomes (30-100 nm in diameter) originate from the multivesicular bodies (MVBs) and are released upon the fusion of MVB with the plasma membrane. In contrast, ectosomes (100-500 nm in diameter) are formed by the outward budding of the plasma membrane (Kalra et al., 2016). Recently, several studies reported that exosomes can contain proteins associated with neurodegenerative diseases and, therefore, may be explored as disease biomarkers (Hill, 2019, Fowler, 2019, You and Ikezu, 2019). However, the role of ectosomes in the pathogenesis of neurodegenerative diseases, and the general effects of EVs in neuronal activity remain largely unknown (Bras et al., 2021).

More recently, another unconventional secretion mechanism known as misfolding-associated protein secretion pathway (MAPS), was described to export misfolded proteins (Xu et al., 2018, Lee et al., 2016). This mechanism uses the ER-associated deubiquitylase USP19 to preferentially export misfolded cytosolic proteins through the recruitment of proteins to the ER surface for deubiquitylation. These cargoes are then encapsulated into late endosomes and secreted to the extracellular space (Lee et al., 2016). After internalization, misfolded proteins may act as seeds to template the misfolding and aggregation of their physiological forms (Jaunmuktane and Brandner, 2020, Sanders et al., 2016, Soto and Pritzkow, 2018).

Here, we developed stable cell lines expressing aSyn, Tau and Htt exon 1 (carrying either 25 or 103 polyglutamines, 25QHtt and 103QHtt, respectively) fused to EGFP in order to afford a systematic comparison of the various proteins. Our results demonstrate that the different disease-related proteins are released, as free forms and in EVs, at different levels. Overall, 25QHtt-EGFP, aSyn-EGFP and EGFP-Tau were found at higher levels in the cell media than 103QHtt-EGFP. We observed similar results when these proteins were expressed in primary cortical neurons or expressed without the EGFP tag, suggesting that the process of release to the extracellular space is mainly dependent on the protein properties. Furthermore, we modelled the occurrence of the proteins in the extracellular space, as when proteins are released from cells, and assessed their effect in the spontaneous firing activity and bursting events of mature primary cortical neurons using multi-electrode arrays (MEA). Monomeric 43QHtt induced discernable alterations in the bursting properties of the cells when compared with 23QHtt or with vehicle-treated cells, suggesting detrimental effects of the polyglutamine expansion on neuronal activity. Interestingly, Tau internalization resulted in increased neuronal activity, with cells exhibiting shorter bursts and higher intra-burst spike frequency, in contrast with only minor alterations observed with aSyn. These results suggest that intrinsic properties of aSyn, Tau or Htt present in the extracellular space, and not necessarily the levels of the proteins, modulate their effect on neuronal activity.

Interestingly, aSyn, Tau and Htt are present at higher levels in ectosomes than in exosomes, without altering the overall proteome of the vesicles. Additionally, the internalization of EVs by microglial or astrocytic cells elicited an increase in the levels of IL-6, IL-1β and TNFα, pro-inflammatory cytokines. Microglial cells also displayed an increase in p62 and LC3 puncta, suggesting the activation of autophagy for digesting the EVs. Finally, neuronal cells also internalized ectosomes and exosomes enriched in aSyn, Tau or Htt and, consequently, exhibited cell bursting irregularities that, overall, correlated with the type of EV used.

Our results indicate that common cellular mechanisms may be used for the transfer of aSyn, Tau and Htt between different cell types. Interestingly, we report that these proteins are handled differently depending on the receptor cell. We posit that the identified similarities and differences between the release and extracellular effects of the three proteins suggest the need for careful consideration of possible targets for therapeutic intervention in different diseases.

## Results

### aSyn, Tau and Htt are released to the extracellular space in different cell models

To evaluate the release of aSyn, Tau and Htt to the extracellular space, we established stable HEK cell lines expressing the different disease-related proteins fused to EGFP (Figure 1). In particular, we expressed two biologically-relevant N-terminal exon 1 Htt fragments with either 25 or 103 polyglutamines (representing wild-type and mutant Htt, 25QHtt-EGFP and 103QHtt-EGFP, respectively) (Figure 1A-B). While 25QHtt-EGFP, aSyn-EGFP and EGFP-Tau expression was mainly diffused in the cytoplasm, 103QHtt-EGFP accumulated in inclusions in the nucleus and throughout the cell (Figure 1A). Importantly, the use of the same cell type expressing the different proteins enabled us to directly compare the effect of the cellular machinery on protein release. We found that aSyn, Tau and Htt were differentially released to the extracellular space (Figure 1B). Cell lysates and conditioned media were assessed by SDS-PAGE, and protein levels were normalized to total protein levels using Memcode (Figure 1B). In particular, the levels of 25QHtt-EGFP in the media were higher than those of 103QHtt-EGFP, aSyn-EGFP or EGFP-Tau (Figure 1B).

**Figure 1.**
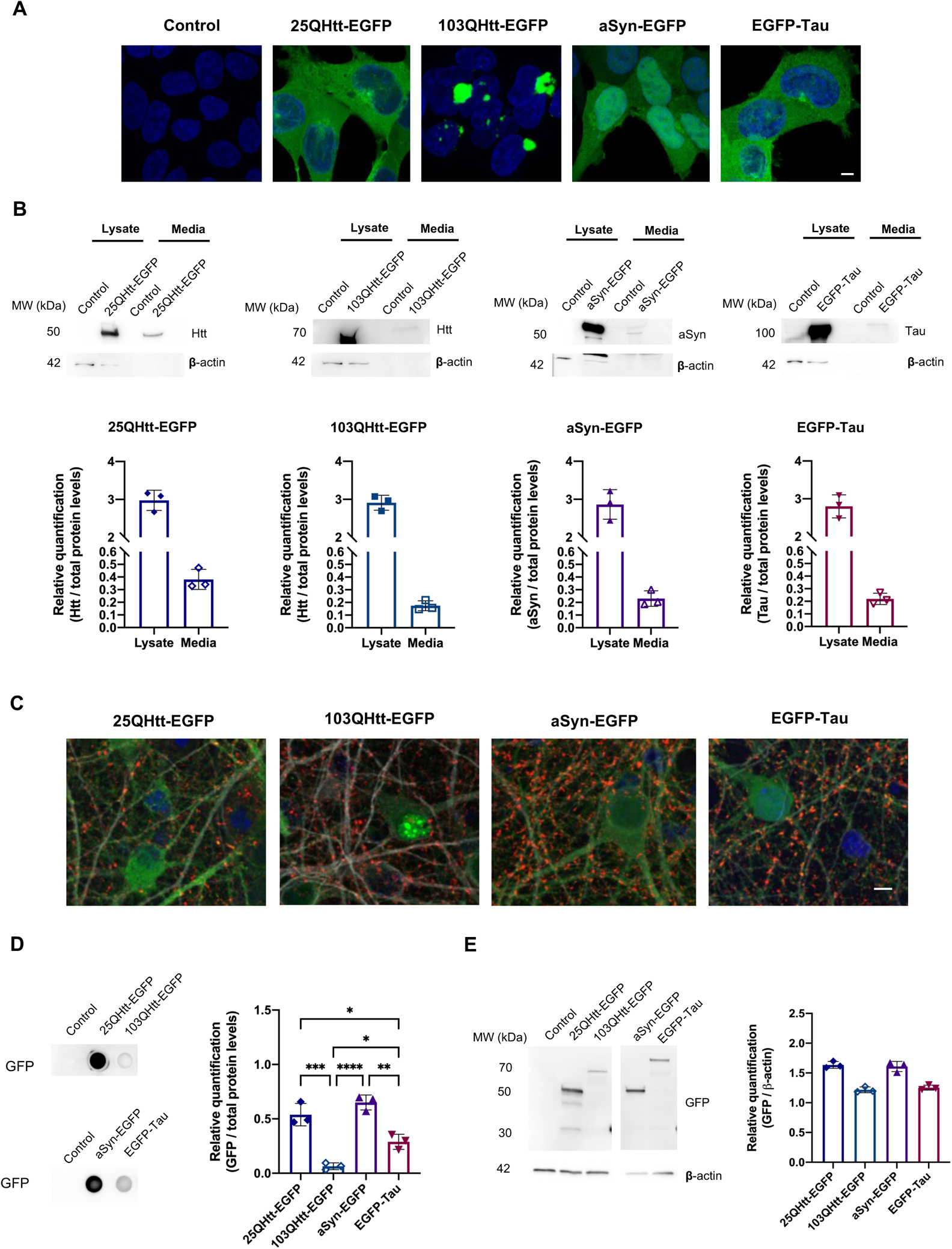
aSyn, Tau and Htt are released to the extracellular space. **(A)** Representative images of HEK cells stably expressing 25QHtt-EGFP, 103QHtt-EGFP, aSyn-EGFP, or EGFP-Tau. Scale bar 5µm. **(B)** Immunoblots showing the protein levels in cell lysates and in the cell media of the different cell lines. Quantifications were normalized to total protein levels using MemCode. **(C)** Representative images of primary cortical neurons infected with lentiviral vectors encoding 25QHtt-EGFP, 103QHtt-EGFP, aSyn-EGFP or EGFP-Tau, and immunostained for synaptophysin (red) and MAP2 (grey). Neurons were infected at DIV14 and cultured until DIV19. They were then fixed for imaging and the media was collected for further analyses. Scale bar 5µm. **(D)** Dot blot analyses of the protein levels released to the cell media of cells expressing the different proteins. Quantifications were normalized to total protein levels using Memcode. **(E)** Immunoblots showing the EGFP levels in whole-cell lysates. Quantifications were normalized to β-actin. Data from at least three independent experiments for each condition. Significant differences were assessed by one-way ANOVA followed by multiple comparisons with significance between groups followed by Bonferroni correction. Differences were considered to be significant for values of p<0.05 and are expressed as mean ± SD, *p<0.05, **p<0.01, ***p<0.001, ****p<0.0001. See also Supplementary Figure 1.

In addition, we evaluated the release of aSyn, Tau and Htt in primary neuronal cultures (Figure 1C-E). Cortical neurons were infected with lentiviral vectors encoding for the various proteins (at DIV14), to ensure homogeneous transduction (Figure 1C). Cell media was collected at DIV19 and applied onto a dot blot system to assess the presence of the different proteins in the extracellular space (Figure 1D). We observed that 25QHtt-EGFP was released at higher levels than 103QHtt-EGFP. Overall, 25QHtt-EGFP and aSyn-EGFP were released at higher levels compared with the other proteins (Figure 1D), but these differences were not simply associated with the expression levels in the cells (Figure 1E).

To rule out that secretion was associated with the presence of the EGFP tag, we expressed untagged aSyn, Tau and Htt proteins in HEK cells (Supplementary Figure 1). Consistently with the previous data, we observed higher levels of 22QHtt, aSyn and Tau in the cell media, and lower levels of 72QHtt (Supplementary Figure 1B). Furthermore, cells did not exhibit signs of toxicity-induced permeabilization, as shown by the lactate dehydrogenase (LDH) cytotoxicity assay (Supplementary Figure 1C - E).

These results demonstrate that aSyn, Tau and Htt are released to the extracellular space at different levels, independently of the cell model used.

### USP19 promotes the secretion of disease-related proteins

Recently, USP19 has been proposed to regulate protein secretion (Lee et al., 2016). To investigate whether MAPS was involved in the release of aSyn, Tau or Htt, we co-expressed USP19 or the catalytic inactive form USP19 C506S with aSyn, Tau or Htt using the HEK stable cell lines we generated above (Figure 2). Interestingly, USP19 significantly increased the secretion of 25QHtt-EGFP after 24 hours (Figure 2A). We also observed a trend towards an increase in the secretion of 103QHtt-EGFP, aSyn-EGFP and EGFP-Tau, but this did not reach statistical significance (Figure 2A). No effects were observed when we co-expressed the catalytic inactive form USP19 C506S, confirming the effects observed were associated with the activity of USP19. Cells did not exhibit signs of toxicity-induced permeabilization (Figure 2B).

**Figure 2.**
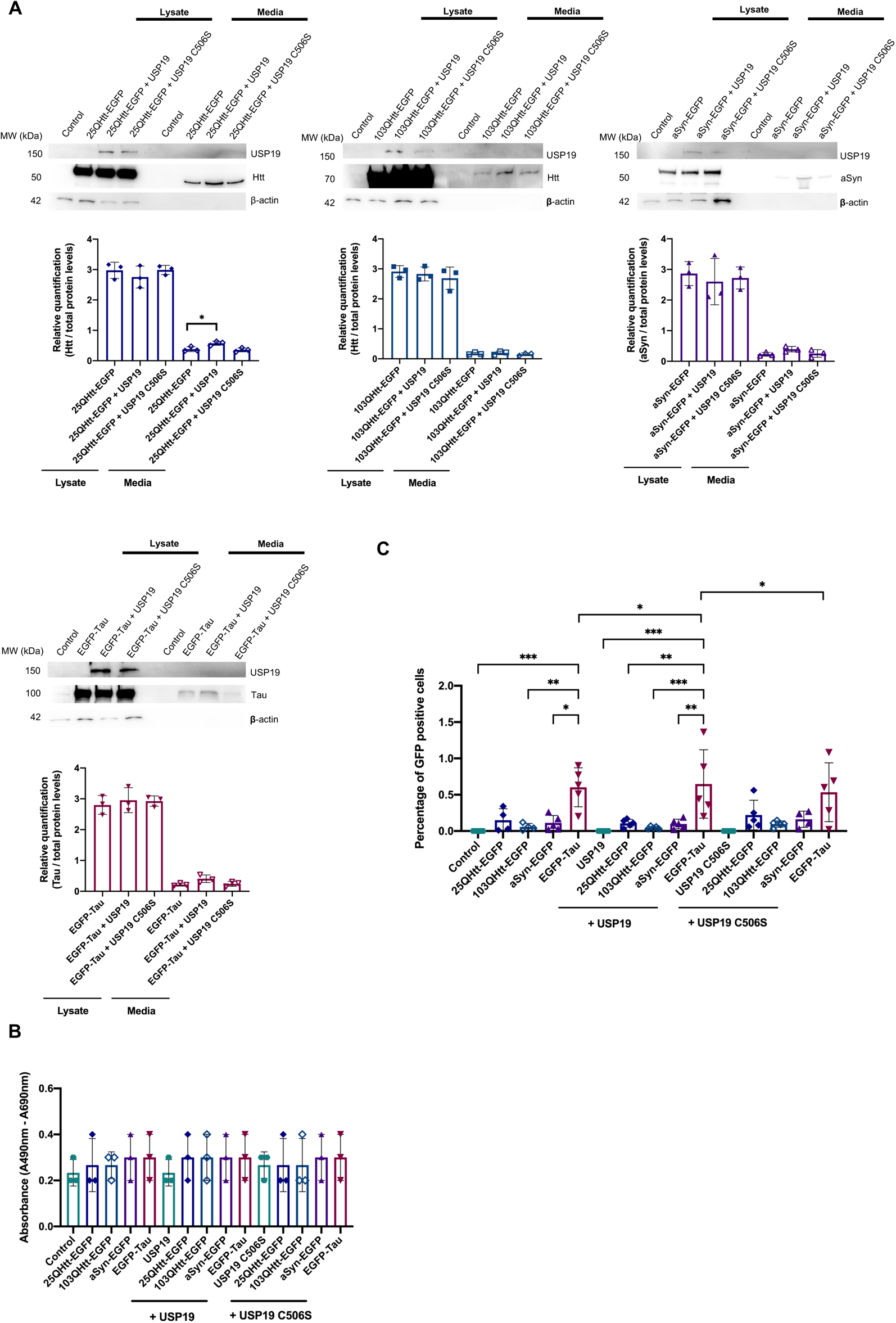
USP19 increases the release of aSyn, Tau or Htt to the extracellular space. **(A)** Immunoblots showing protein levels in cell lysates and released to the cell media of cells expressing the different proteins. HEK cell stably expressing 25QHtt-EGFP, 103QHtt-EGFP, aSyn-EGFP and EGFP-Tau were transfected with USP19 or with the catalytic inactive form USP19 C506S. Quantifications were normalized to total protein levels using MemCode. **(B)** LDH measurements confirm the absence of cell toxicity and cell death in the experiments. **(C)** Tau is more strongly internalized by naïve cells. Percentage of EGFP positive cells after incubation with media from cells co-expressing 25QHtt-EGFP, 103QHtt-EGFP, aSyn-EGFP or EGFP-Tau together with USP19 or USP19 C506S for 24 hours. Cell counting was performed using flow cytometry. Data from at least three independent experiments for each condition. Significant differences were assessed by one-way ANOVA followed by multiple comparisons with significance between groups corrected by Bonferroni procedure. Differences were considered to be significant for values of p<0.05 and are expressed as mean ± SD, *p<0.05, **p<0.01, ***p<0.001, ****p<0.0001.

Next, we tested whether the secreted proteins could be internalized by naive cells. For this, the cell media from the different stable cell lines was collected after 24 hours, shortly centrifuged to deplete possible floating cells present in the media, and then added to naïve HEK cells for 72 hours. The number of EGFP-positive cells was analyzed using flow cytometry, and we found a higher percentage of EGFP-Tau positive cells compared with the other conditions (Figure 2A, 2C). Strikingly, EGFP-Tau was not the protein released at higher levels to the cell media (Figure 2A). Interestingly, we also observed that incubation of naïve cells with media from cells co-expressing EGFP-Tau and USP19 resulted in a higher percentage of positive cells, indicating higher internalization levels when compared with the other proteins (Figure 2C).

Together, these results indicate that the levels of protein secretion are not directly correlated with the levels of internalization by receiving cells and may, instead, depend on intrinsic properties of the proteins and on the pathways involved in protein uptake.

### Disease-related proteins are present at higher levels in ectosomes

Previous research indicates that aSyn, Tau and Htt can be secreted in association with EVs (You and Ikezu, 2019, Beatriz et al., 2021). To assess the contribution of ectosomes and exosomes towards the transfer of aSyn, Tau and Htt, we used an optimized differential ultracentrifugation protocol to purify these EVs from the cell media of HEK cells stably expressing the different proteins (Figure 3, Supplementary Figure 2) (Bras et al., 2021). Staining of ectosomal and exosomal fractions showed a similar total protein profile that, as expected, was distinct from that of the whole-cell lysate (Supplementary Figure 2A). Electron microscopy (EM) imaging confirmed the greater diameter of ectosomes in comparison to exosomes, and their characteristic cup-shape derived from the ultracentrifugation protocol (Supplementary Figure 2B). In addition, the size distribution and concentration of the two EV types was further validated using Nanosight (Supplementary Figure 2C). While ectosomes presented a diameter of ∼140nm, the diameter of exosomes was ∼60nm (Supplementary Figure 2C). Conventional exosomal protein markers such as alix, flotillin-1 or TSG101 were clearly enriched in the exosomal fraction, whereas the ectosomal fraction was enriched in annexin-A2 and annexin-A5 (evaluated using immunoblot and mass spectrometry) (Figure 3A, Supplementary Figure 2D). The ER and Golgi markers calnexin and GM130, respectively, were not detected, confirming the high purity of the isolated EVs (Supplementary Figure 2D).

**Figure 3.**
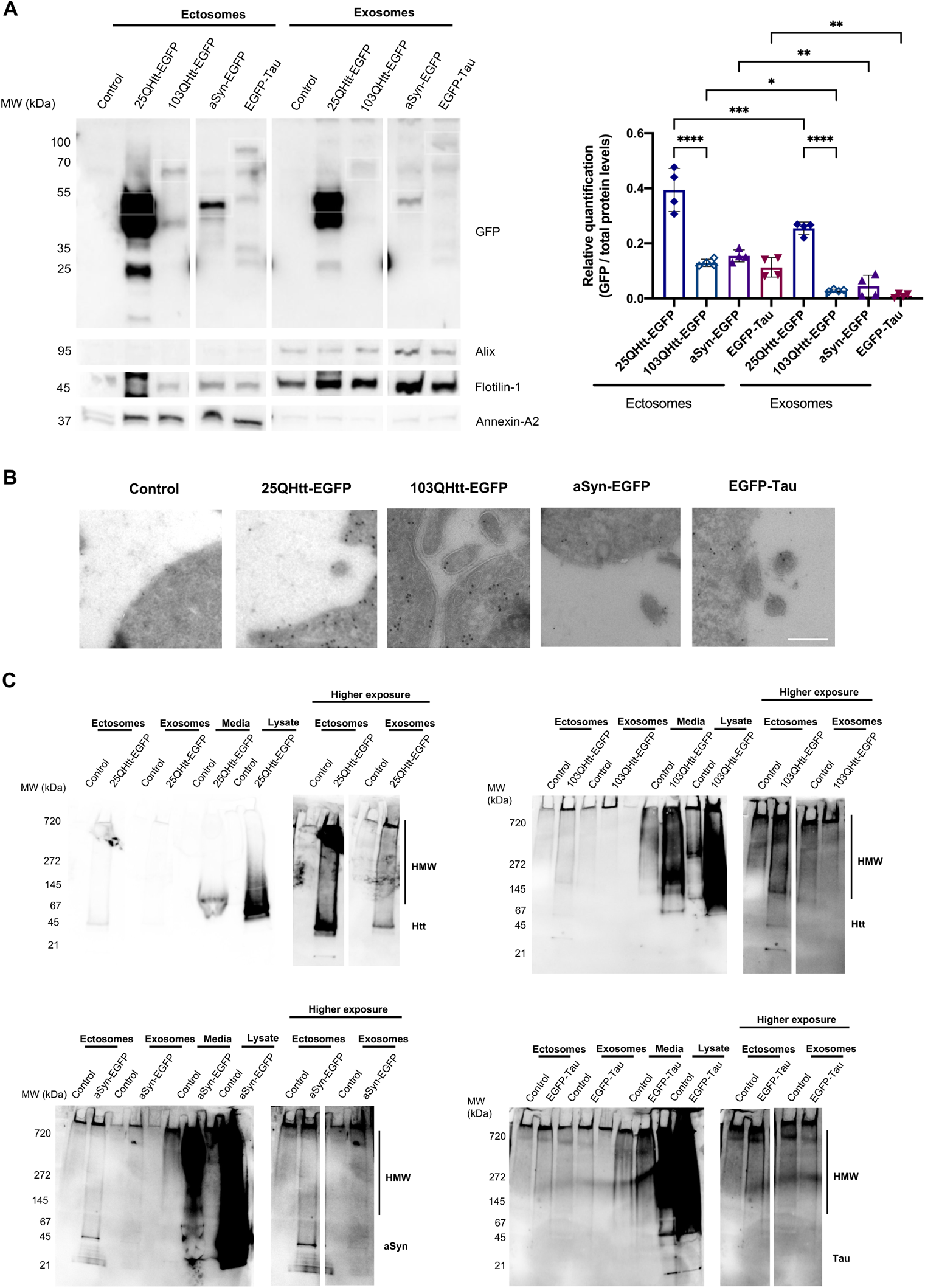
Disease-related proteins are enriched in ectosomes. **(A)** Immunoblots of ectosomal and exosomal fractions purified from the media of HEK cells stably expressing 25QHtt-EGFP, 103QHtt-EGFP, aSyn-EGFP or EGFP-Tau for 24 hours. Equal amounts of protein were separated on SDS-PAGE gels, and membranes were incubated with the indicated antibodies. Protein levels were normalized to total protein levels using Memcode. **(B)** Representative immunoelectron microscopy images of HEK cells stably expressing 25QHtt-EGFP, 103QHtt-EGFP, aSyn-EGFP or EGFP-Tau immunolabelled for EGFP, as a common marker (scale bar 200 nm). **(C)** Ectosomes and exosomes containing disease-related proteins have high molecular weight (HMW) species. Ectosomal and exosomal fractions, cell media and whole-cell lysates of HEK cells stably expressing 25QHtt-EGFP, 103QHtt-EGFP, aSyn-EGFP or EGFP-Tau. Equal amounts of the proteins were separated on native gels, and membranes were incubated with the indicated antibodies. Data from at least three independent experiments for each condition. Significant differences were assessed by one-way ANOVA followed by multiple comparisons with significance between groups corrected by Bonferroni procedure. Differences were considered to be significant for values of p<0.05 and are expressed as mean ± SD, *p<0.05, **p<0.01, ***p<0.001, ****p<0.0001. See also Supplementary Figures 2-3.

Interestingly, aSyn, Tau and Htt were detected at higher levels in ectosomes than in exosomes (EGFP levels were normalized to the total protein levels in the immunoblot using MemCode) (Figure 3A). These results were further confirmed using antibodies specific for aSyn, Tau or Htt, and by mass spectrometry analyses (Supplementary Figure 3). We also detected S129 phosphorylation of aSyn, a posttranslational modification (PTM) typically associated with pathology, in the lysates of aSyn-EGFP expressing cells, but not in the EV fractions (Supplementary Figure 3A). Surprisingly, ectosomes and exosomes containing 25QHtt-EGFP, 103QHtt-EGFP, aSyn-EGFP or EGFP-Tau presented similar proteomic signatures when compared with EVs purified from control cells (Supplementary Figure 3B-C).

Immuno-EM experiments demonstrated the presence of the different disease-related proteins in the cytoplasm, as expected, but also near to the plasma membrane, implying their availability to be incorporated in ectosomes (Figure 3B). Next, to assess the biochemical state of aSyn, Tau and Htt in the cell media and in EVs, the different samples were applied onto a native gel (Figure 3C). Overall, cell media presented greater levels of high molecular species when compared with the EV fractions, possibly due to the higher levels of 25QHtt-EGFP, 103QHtt-EGFP, aSyn-EGFP or EGFP-Tau present in the cell media. Furthermore, ectosomes containing the disease-related proteins presented a stronger smear when compared with exosomes.

These results highlight the prominent role ectosomes, and not only exosomes, may play in the release of disease-related proteins to the extracellular space.

### EVs containing aSyn, Tau and Htt are internalized by glial cells and induce the production of pro-inflammatory cytokines

Microglial and astrocytic cells produce neuroinflammatory cytokines and appear to be involved in the spreading in neurodegenerative diseases (Budnik et al., 2016, Coleman and Hill, 2015, Franklin et al., 2021). Therefore, we then asked whether ectosomes and exosomes containing aSyn, Tau and Htt were internalized by different brain cell types, and whether the disease-associated protein present in the vesicles would alter the responses elicited (Figure 4).

**Figure 4.**
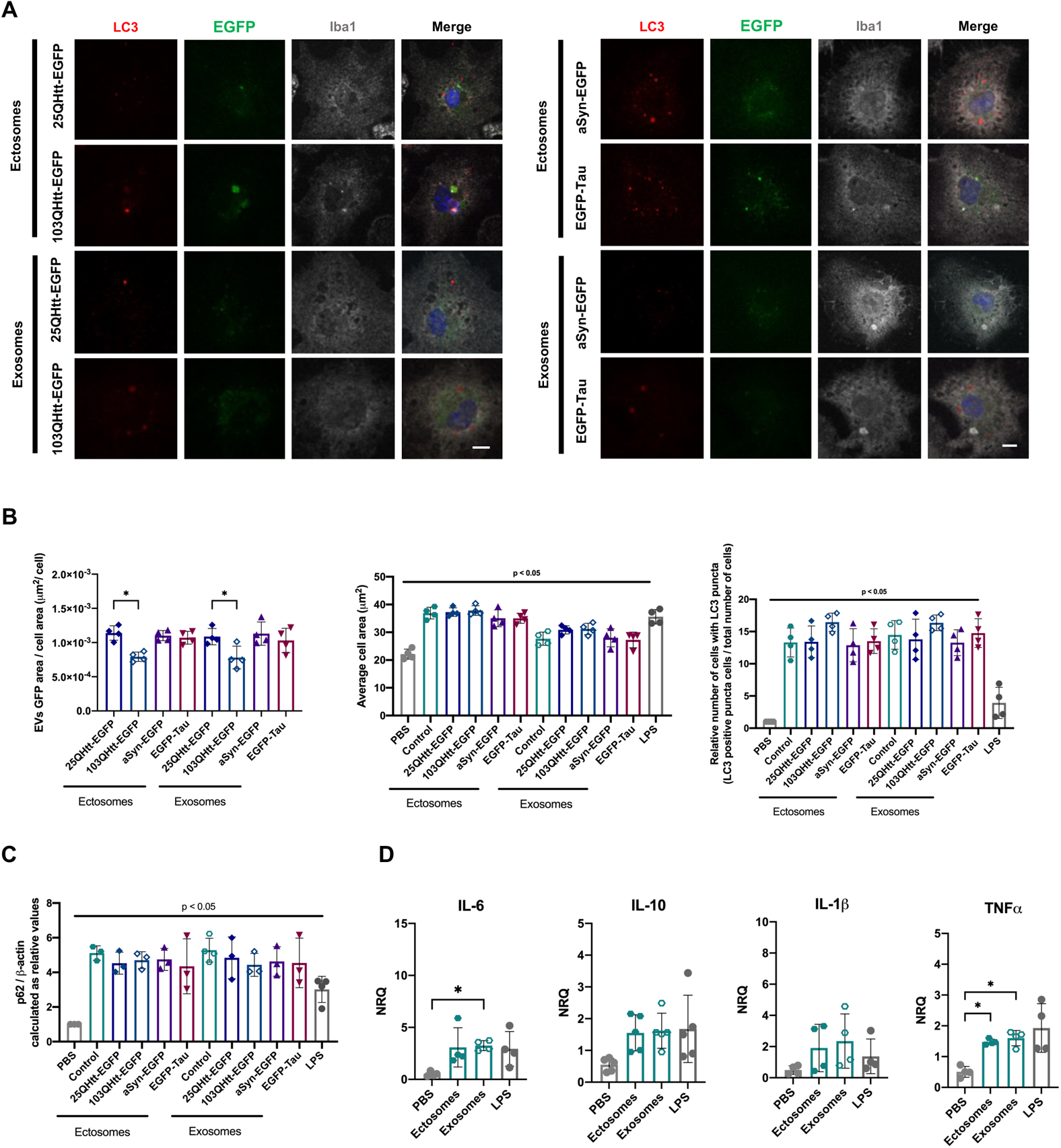
Ectosomes and exosomes containing disease-related proteins are internalized by microglial cells. **(A)** Ectosomes and exosomes from cells expressing 25QHtt-EGFP, 103QHtt-EGFP, aSyn-EGFP or EGFP-Tau were applied to microglial cultures at a concentration of 10*μ*g/mL for 24 hours. Cells were immunoassayed for LC3 (red) and Iba1 (grey). Scale bar 10 µm. **(B)** EV internalization was evaluated through imaging analysis by measuring EGFP signal and cell area. Microglial cells treated with EVs show an increase in average cell area and number of cells containing LC3-positive puncta. **(C)** Increase in p62 levels after 24hours of EV treatment. Quantifications were normalized to β-actin levels. **(D)** EV treatment resulted in the activation of the pro-inflammatory markers IL-6 and TNFα in microglia cells after 24hours. Data from at least three independent experiments for each condition. Significant differences were assessed by one-way ANOVA followed by multiple comparisons with significance between groups corrected by Bonferroni procedure. Differences were considered to be significant for values of p<0.05 and are expressed as mean ± SD, *p<0.05, **p<0.01. See also Supplementary Figures 4-5.

Primary microglial cells were treated with 10*μ*g/mL of ectosomes and exosomes purified from the media of cells expressing 25QHtt-EGFP, 103QHtt-EGFP, aSyn-EGFP or EGFP-Tau (Figure 4, Supplementary Figure 4-5). Furthermore, cells were also exposed to the bacterial endotoxin lipopolysaccharide (LPS) as a pro-inflammatory stimulus, as a control (Lively and Schlichter, 2018). After 24 hours of treatment, EGFP–labeled ectosomes and exosomes were found to colocalize with the microglial marker Iba1 in the cell cytoplasm, indicating EVs were internalized (Figure 4A, Supplementary Figure 4A). Interestingly, EVs containing 103QHtt-EGFP were less internalized, or degraded more rapidly, when compared with EVs containing 25QHtt-EGFP (Figure 4B). Overall, the internalization ratio was similar for all ectosomes and exosomes tested (Figure 4B). Labelling of EVs with the thiol-based dye Alexa Fluor 633 C5-maleimide further confirmed the previous results (Supplementary Figure 4B-C) (Roberts-Dalton et al., 2017). EV internalization resulted in microglia activation, as demonstrated by an increase in cell area and elevated levels of the pro-inflammatory cytokines IL-6 and TNFα (Figure 5B-D). These results were independent of the presence of aSyn, Tau or Htt in the vesicles (Supplementary Figure 5). Furthermore, EV uptake resulted in an increase in autophagosomes in the cytoplasm (increase in LC3 puncta) and higher p62 levels, indicating autophagy activation (Hung et al., 2009) (Figure 4B-C). No differences in the levels of iNOS, Iba1, APG5L/ ATG5 or LC3 were observed, and no toxicity was observed after the treatment (Supplementary Figure 5).

**Figure 5.**
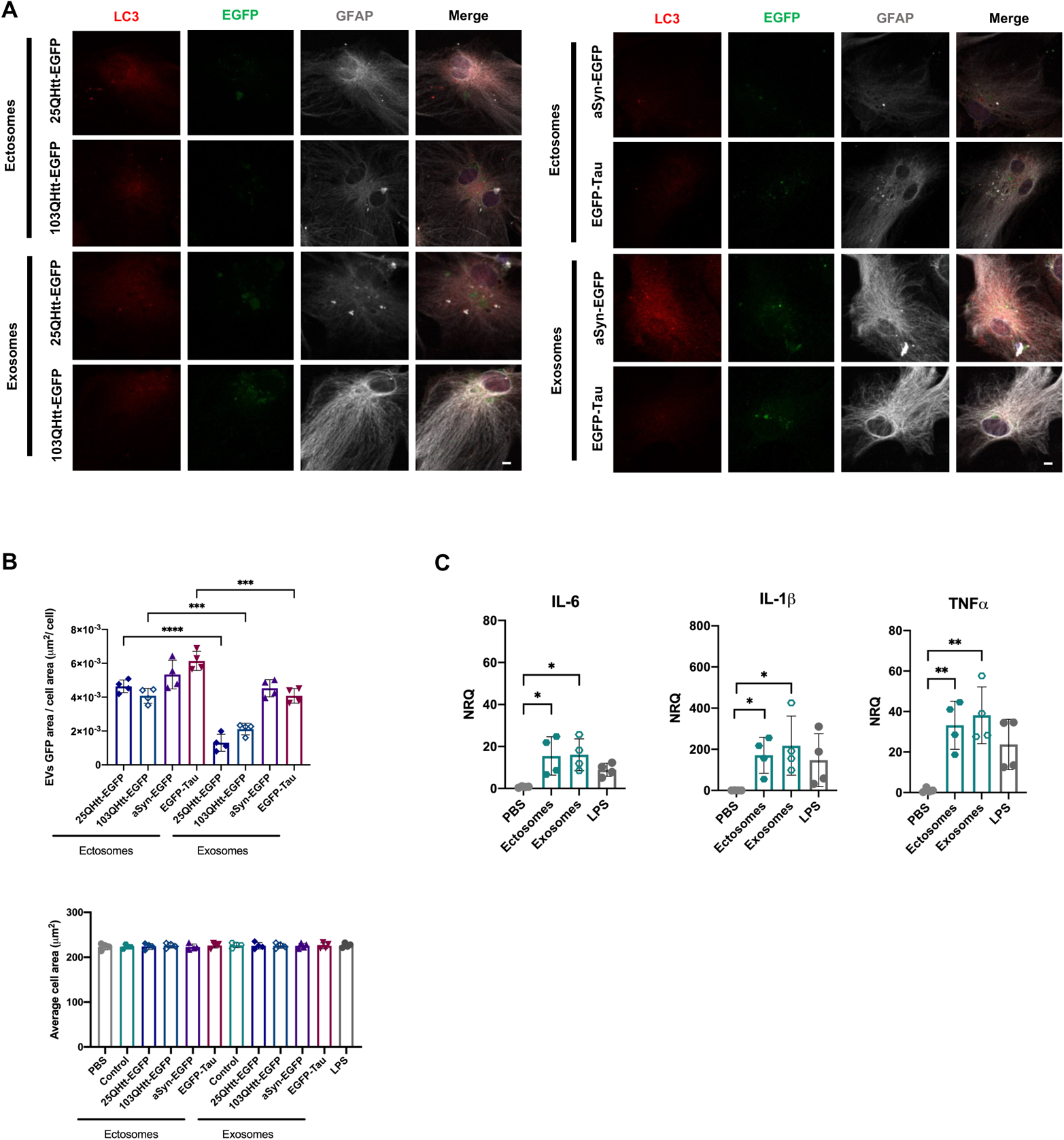
Ectosomes and exosomes containing disease-related proteins are internalized by astrocytic cells. **(A)** Ectosomes and exosomes from cells expressing 25QHtt-EGFP, 103QHtt-EGFP, aSyn-EGFP or EGFP-Tau were applied to astrocytic cultures at a concentration of 20*μ*g/mL for 24 hours. Cells were immunoassayed for LC3 (red) and GFAP (grey). Scale bar 10 µm. **(B)** Astrocytic cells treated with EVs do not change average cell area. EVs internalization levels were evaluated through imaging analysis by measuring EGFP signal and cell area. **(C)** EV treatment resulted in the activation of the pro-inflammatory markers IL-6, IL-β and TNFα in astrocytic cells after 24 hours. Data from at least three independent experiments for each condition. Significant differences were assessed by one-way ANOVA followed by multiple comparisons with significance between groups corrected by Bonferroni procedure. Differences were considered to be significant for values of p<0.05 and are expressed as mean ± SD, *p<0.05, **p<0.01, ***p<0.001. See also Supplementary Figures 6-7.

To further explore the effect of EV internalization in glial cells, primary astrocytes were treated with 20*μ*g/mL of ectosomes and exosomes purified from cells expressing 25QHtt-EGFP, 103QHtt-EGFP, aSyn-EGFP or EGFP-Tau (Figure 5, Supplementary Figure 6-7). After 24 hours of treatment, we observed the colocalization of the astrocytic marker GFAP with EGFP–labeled ectosomes and exosomes, confirming internalization (Figure 5A, Supplementary Figure 6A). Interestingly, after internalization, the EGFP signal of the EVs was surrounded by LC3 signal, suggesting a possible engulfment and degradation of EVs in the cells (Figure 5A, Supplementary Figure 6A-C). Quantification of the internalization ratio indicated greater engulfment of ectosomes than exosomes, and these results were further confirmed using dye-labelled EVs (Figure 5B, Supplementary Figure 6D). Internalization of ectosomes and exosomes caused an increase in pro-inflammatory cytokines, including IL-6, IL-1β and TNFα, without changing the average cell area (Figure 5B-C). These results were independent of the presence of aSyn, Tau or Htt in the vesicles (Supplementary Figure 7). No significant differences in the levels of iNOS, APG5L/ ATG5, p62, GFAP or LC3 were observed after EV treatment, and no cytotoxicity was detected (Supplementary Figure 7).

These results indicate that different types of EVs can be internalized in microglial and astrocytic cells, independently of the presence of aSyn, Tau and Htt in the vesicles. Furthermore, EV internalization elicited similar responses, with an increase in inflammatory markers and autophagy activation.

### Treatment with EVs containing aSyn, Tau and Htt modifies spontaneous neuronal activity in cortical neurons

Since EVs are actively released to the extracellular space by different types of brain cells, we hypothesized that spontaneous neuronal activity might be influenced by the internalization of ectosomes and/or exosomes in neuronal cells (Bras et al., 2021). To test this, we treated primary cortical neurons with 20*μ*g/mL of ectosomes and exosomes purified from cells expressing 25QHtt-EGFP, 103QHtt-EGFP, aSyn-EGFP or EGFP-Tau, at DIV14, for 24 hours to allow internalization of EVs (Figure 6, Supplementary Figure 8). We observed neuronal cells internalized ectosomes and exosomes at similar levels, and that the EV signal was surrounded by LC3 staining, suggesting the targeting of the EVs for degradation after uptake (Figure 6A-C, Supplementary Figure 8A). The internalization of EVs did not alter the average cell area or caused toxicity and it did not induce significant differences in synaptic and autophagic protein levels (Figure 6D, Supplementary Figure 8C-D).

**Figure 6.**
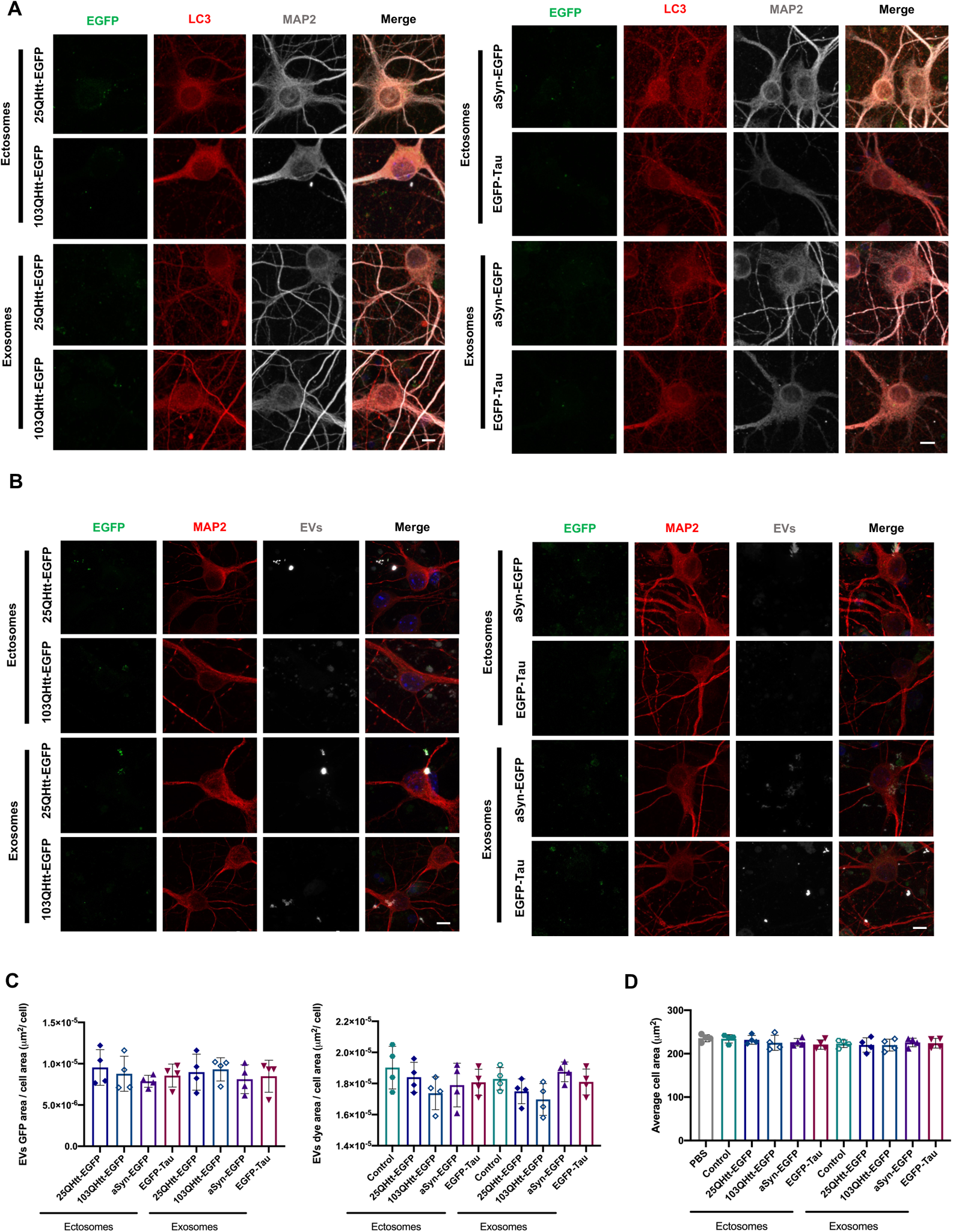
Ectosomes and exosomes containing disease-related proteins are internalized by primary cortical neurons. **(A)** Ectosomes and exosomes from cells expressing 25QHtt-EGFP, 103QHtt-EGFP, aSyn-EGFP or EGFP-Tau were applied to primary cortical neurons 20*μ*g/mL for 24 hours. Cells were immunoassayed for LC3 (red) and MAP2 (grey). Scale bar 5 µm. **(B)** Ectosomes and exosomes containing 25QHtt-EGFP, 103QHtt-EGFP, aSyn-EGFP or EGFP-Tau were labelled with Alexa Fluor 633 C5-maleimide (gray) and applied to cell cultures at a concentration of 20*μ*g/mL for 24 hours. Cells were immunostained for MAP2 (red). Scale bar 5µm. **(C)** EV internalization was evaluated through imaging analysis by measuring EGFP signal, dye area, and cell area. **(D)** Neuronal cells treated with EVs do not change average cell area. Data from at least three independent experiments for each condition. Significant differences were assessed by one-way ANOVA followed by multiple comparisons with significance between groups corrected by Bonferroni procedure. Differences were considered to be significant for values of p<0.05 and are expressed as mean ± SD. See also Supplementary Figures 8.

Next, we assessed the effects induced by the EVs on neuronal activity using multi-electrode arrays (MEAs) (Bras et al., 2021). Primary cortical neurons were cultured in MEA chambers and treated with 20*μ*g/mL of EVs at DIV14. Firing activity was recorded 24 hours after treatment with ectosomes or exosomes purified from cells expressing 25QHtt-EGFP, 103QHtt-EGFP, aSyn-EGFP or EGFP-Tau (Figure 7). Representative raster plots showed the individual firing activity and bursts events in neuronal cultures for each treatment condition (Figure 7A). We observed a reduction in the mean firing rate after EV internalization, and a decrease in the average spike amplitude for the neurons treated with exosomes (Figure 7B). The assessment of bursting activity parameters showed that, upon treatment with EVs, spike bursts were more irregular and with longer duration (Figure 7C). Overall, exosome internalization resulted in longer inter-burst intervals with reduced intra-burst spiking frequency, and a reduced percentage of spikes within bursts, in contrast to what we observed in neurons treated with ectosomes (Figure 7C). Furthermore, these alterations were more strongly correlated with the EV subtype, and not with the presence of aSyn, Tau or Htt.

**Figure 7.**
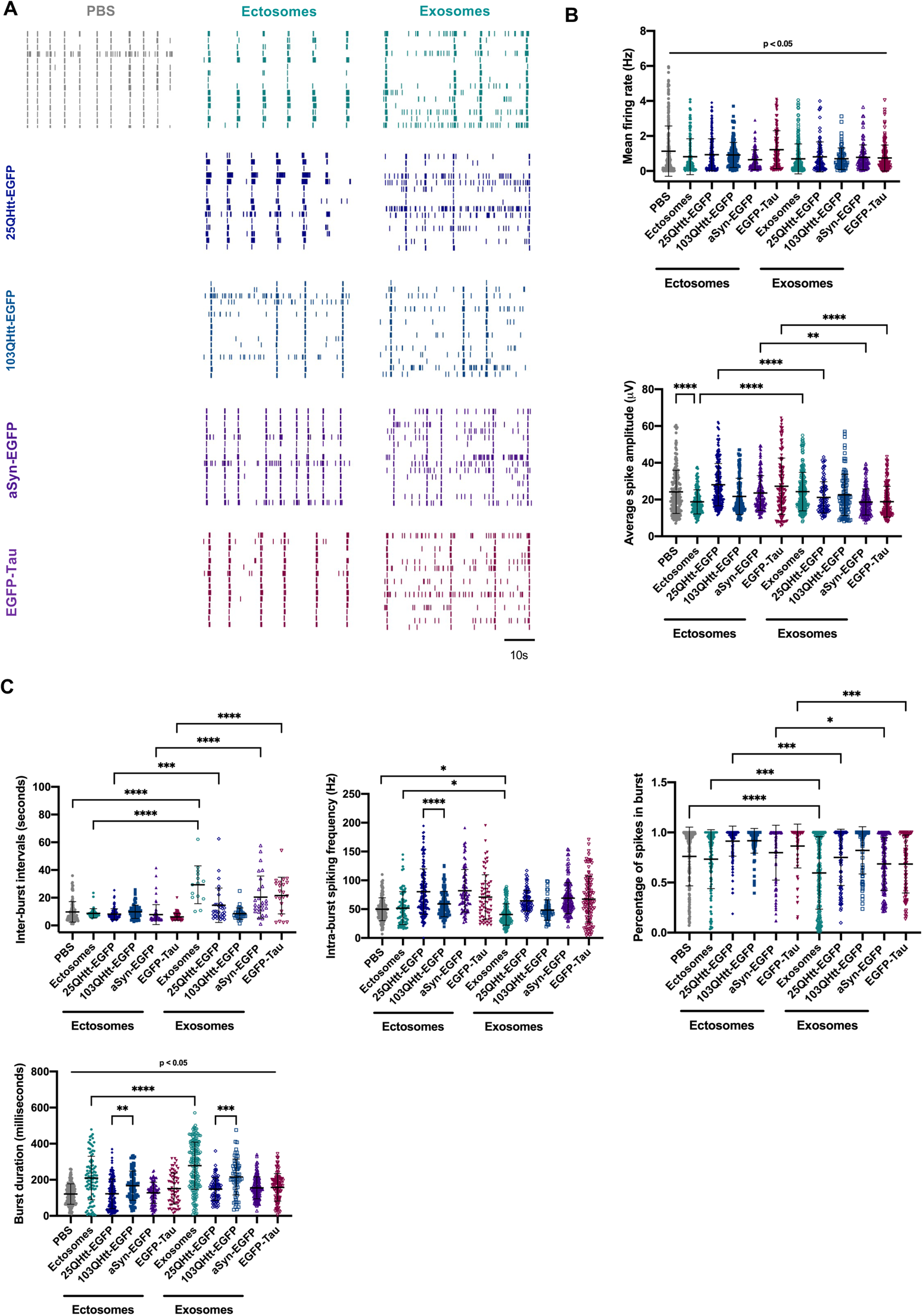
EVs containing disease-related proteins modulate spontaneous activity in primary cortical neurons. **(A)** Representative raster plots of the spontaneous firing activities recorded from cortical neurons after incubation with 20µg/mL ectosomes and exosomes control or containing aSyn-EGFP, EGFP-Tau, 25QHtt-EGFP and 103QHtt-EGFP at DIV14 for 24 hours, recorded using 60-electrode MEAs. In every block, each row represents one single cell (15 cells shown) and each vertical line represents a single spike obtained on DIV15 [scale bar represents 10 seconds (s)]. **(B)** Quantification of the mean firing rate and average spike amplitude from primary cortical neurons incubated with PBS or disease-related proteins. **(C)** Bursting properties of the cortical neurons treated with PBS or EVs (inter-burst intervals, intra-burst spiking frequency, percentage of spikes in bursts and burst duration). Data from at least three independent experiments for each condition. Significant differences were assessed by one-way ANOVA followed by multiple comparisons with significance between groups corrected by Bonferroni procedure. Differences were considered to be significant for values of p<0.05 and are expressed as mean ± SD, *p<0.05, **p<0.01, ***p<0.001, ****p<0.0001.

Altogether, our results indicate that, after exposure to EVs, the firing of cultured neurons is more irregular, highlighting the potential of ectosomes and exosomes in modifying important aspects of spontaneous neuronal activity.

### Neuronal network activity is differentially modulated by the internalization of monomeric aSyn, Tau or Htt

Our results demonstrate the release of wild-type and pathogenic forms of disease-related proteins to the extracellular space. Although these proteins are taken up by cells, it is still not known whether and how they modulate neuronal activity (Brahic et al., 2016, Monsellier et al., 2016, Ruiz-Arlandis et al., 2016, Pieri et al., 2012, Evans et al., 2018, Frost et al., 2009, Wu et al., 2013). To address the functional effects of free forms of aSyn, Tau and Htt, we treated primary cortical neurons with 100nM of monomeric aSyn, Tau, 23QHtt or 43QHtt for 5 days, starting at DIV14 (this concentration was selected as it demonstrated no detrimental effects, which would be incompatible with long-term observations) (Figure 8) (Bras et al., 2021).

**Figure 8.**
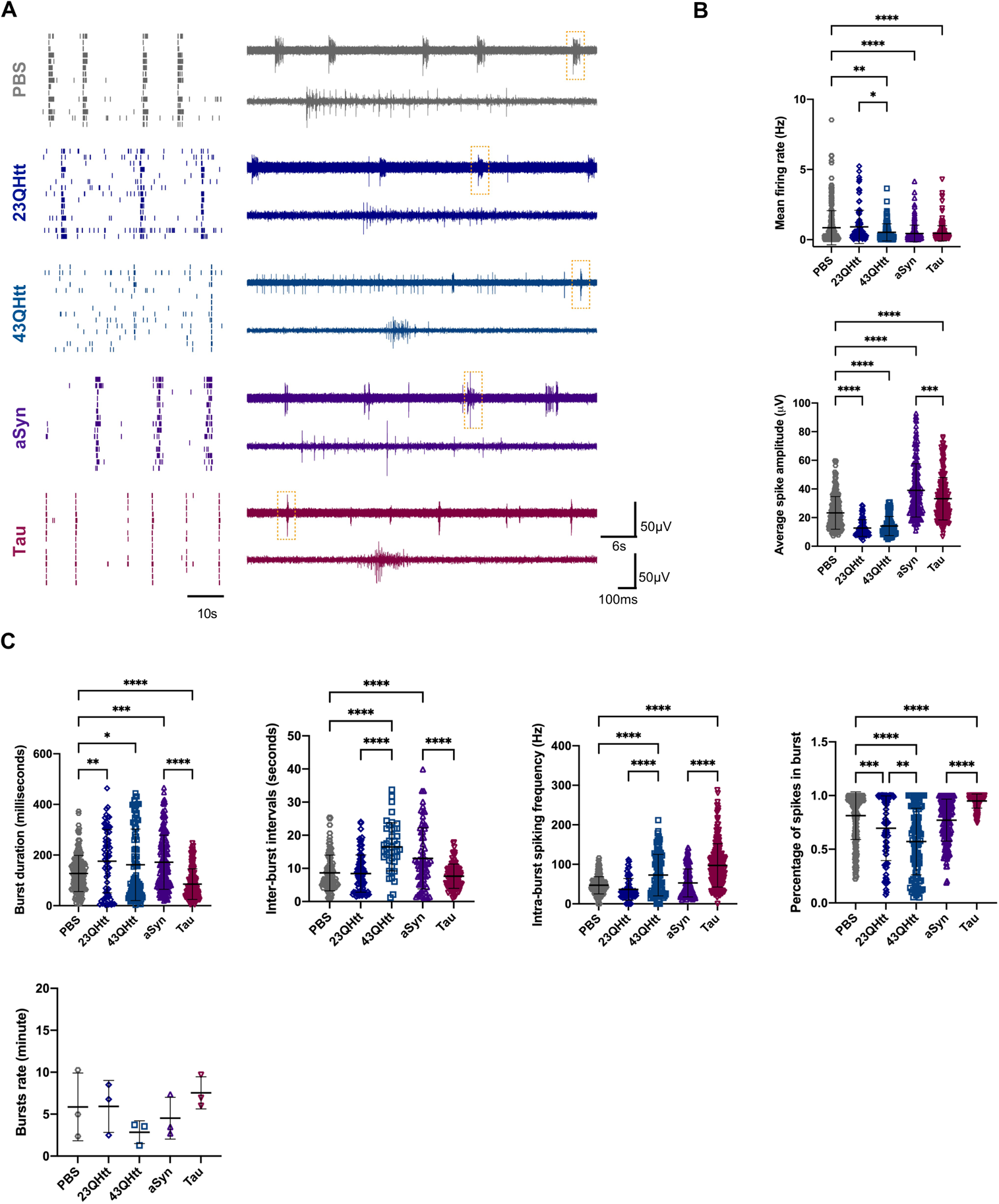
Free monomeric disease-related proteins modulate neuronal activity in primary cortical neurons. **(A)** On the left, representative raster plots of the spontaneous firing activities recorded from cortical neurons after incubation with 100nM of recombinant protein at DIV14 for 5 days, recorded using 60-electrode MEAs. In every block, each row represents one single cell (15 cells shown) and each vertical line represents a single spike obtained on DIV19 [scale bar represents 10 seconds (s)]. On the right, representative voltage traces showing the typical firing activity and bursts events in neuronal cultures treated with PBS, monomeric 23Qhtt, 43QHtt, aSyn and Tau (upper traces, scale bars represent 60µV and 6s). Closeups of the dashed boxes represent the spikes occurring within a burst ([lower traces, scale bars represent 60µV and 100 milliseconds (ms)]. **(B)** Quantification of the mean firing rate and average spike amplitude from primary cortical neurons incubated with PBS or disease-related proteins. **(C)** Bursting properties of the cortical neurons treated with PBS or the disease-related proteins (burst duration, inter-burst intervals, intra-burst spiking frequency, percentage of spikes in bursts and burst rate). Data from at least three independent experiments for each condition. Significant differences were assessed by one-way ANOVA followed by multiple comparisons with significance between groups corrected by Bonferroni procedure. Differences were considered to be significant for values of p<0.05 and are expressed as mean ± SD, *p<0.05, **p<0.01, ***p<0.001, ****p<0.0001.

Representative raster plots and voltage traces show the firing activity and burst events in neuronal cultures treated with PBS (as a negative control), 23QHtt, 43QHtt, aSyn or full-length Tau (Figure 8A). Overall, neurons treated with monomeric proteins showed a reduction in the mean firing rate. In addition, neurons treated with Htt exhibited a decrease in the average spike amplitude (Figure 8B). Remarkably, treatment of neuronal cultures with monomeric aSyn or Tau resulted in an increase in the average spike amplitude (Figure 8B). Furthermore, we observed that Tau induced condensed and more intense spike bursting, as demonstrated by bursts with shorter duration and higher intra-burst spike frequency. Also, the percentage of spikes in each burst increased, suggesting that neurons fire in a more regular and synchronized manner (Figure 8C). Cells treated with recombinant aSyn exhibited slight alterations in their activity, with longer burst duration and larger intervals between them. Treatment with monomeric 23QHtt or 43QHtt resulted in bursts with longer intervals and a reduction in the number of spikes within the burst when compared with the control. Interestingly, 43QHtt induced a stronger alteration in the spontaneous neuronal activity with a slight reduction in burst rate and a decrease in the percentage of spikes per burst when compared with 23QHtt or PBS-treated cells. In addition, neurons displayed longer inter-burst intervals and higher intra-spike frequency, which was possibly correlated with toxic effects induced by the polyglutamine expansion.

Altogether, our results demonstrate that different disease-related proteins induce specific effects on spontaneous neuronal activity.

## Discussion

Several neurodegenerative disease-related proteins appear to be transferred from cell-to-cell, contributing for the spreading of pathology and disease progression (Peng et al., 2020, Bras and Outeiro, 2021, Alpaugh and Cicchetti, 2021). However, the basic molecular mechanisms involved in the release of proteins which are, oftentimes, not typical secretory proteins, are still unclear. Likewise, the effect of such proteins once they are in the extracellular milieu, is also unclear. Therefore, deciphering the pathways through which brain cells release, sense, and respond to the extracellular presence of normal and pathological forms of disease-associated proteins is essential. Here, we conducted a systematic comparison of the basic molecular mechanisms involved in the release of three proteins associated with distinct neurodegenerative disorders, aSyn, Tau, and Htt, and of general cell-autonomous responses in different brain cell types. Since these proteins accumulate in different types of brain cells, forming distinct proteinaceous inclusions, and spreading through different neuronal circuits, it is important to establish differences and similarities in the ways they are handled in order to identify specific therapeutic targets for each disease. In our study, exploiting simple, yet tractable, cell systems, we found that aSyn, Tau and Htt are transferred between cells at different levels, but using overlapping cellular pathways. Importantly, we report that the release of these proteins in a “free form”, or in extracellular vesicles, elicits different molecular processes in neighboring cells.

By taking advantage of stable cell lines expressing aSyn, Tau, 25QHtt and 103QHtt fused to EGFP, as a common denominator, we were able to study and compare different molecular mechanisms involved in the release, and uptake of the various in receptor cells. We demonstrate that 25QHtt-EGFP, aSyn-EGFP and EGFP-Tau are released to the extracellular space at higher levels than 103QHtt-EGFP. Importantly, the presence of the EGFP tag did not alter the release patterns of aSyn, Tau and Htt.

Our study further implicates the MAPS pathway in the release of a fraction of aSyn, Tau or Htt. In agreement with previous studies, USP19 expression led to a slight increase in the release of aSyn, Tau and Htt (Xu et al., 2018, Lee et al., 2016). Interestingly, USP19 promoted a significant increase in the secretion of 25QHtt-EGFP, suggesting a relevant role for this pathway in HD, as it was previously described to interact and regulate mutant Htt protein levels and promote its aggregation (He et al., 2016, He et al., 2017).

After release, aSyn, Tau or Htt were found to be internalized by naïve receptor cells. In particular, we observed a significant increase in the percentage of EGFP-positive cells after incubation of naïve cells with media collected from cells expressing EGFP-Tau. Although 25QHtt-EGFP and aSyn-EGFP were more abundant in the cell media than EGFP-Tau, we observed more internalization of this protein in cells. These results suggest that Tau might be more easily, or more rapidly, internalized by cells, or that different pathways might be involved in its uptake when compared with the other proteins. Strikingly, this observation also demonstrates that protein internalization is not a process solely dependent of the protein levels in the exterior space but determined by the type of protein and mechanisms involved in the cellular uptake.

In addition to the release of proteins in free form, several studies have also reported the secretion of disease-related proteins in ectosomes and exosomes (You and Ikezu, 2019, Tang, 2018). Exosomes are the most extensively studied type of EVs, and are also implicated in the secretion of pathological proteins (Beatriz et al., 2021). However, Tau was also found to be present in ectosomes extracted from culture media from cell models and human cerebrospinal fluid (Dujardin et al., 2014, Spitzer et al., 2019), highlighting the relevance of this EV type, and need for further research to address their role in neurodegenerative diseases. In our study, we found that ectosomes purified from the cell media of stable cell lines contained higher levels of aSyn, Tau and Htt than exosomes. Interestingly, we observed that aSyn, Tau and Htt are present near the plasma membrane, in agreement with their possible incorporation in ectosomes, and release via passive diffusion. Traditionally, ectosome characterization has been challenging due to the lack of specific protein markers. In our study, we highlight the enrichment of annexin-A2 in ectosomes, suggesting this protein as a specific marker for this EV type, as previously described (Bras et al., 2021).

We also found that incorporation of aSyn, Tau or Htt in EVs did not change the normal vesicle protein composition, suggesting these vesicles are not deregulated when they transport the disease-associated proteins we tested. However, future research will be necessary to determine whether the content of other biomolecules, such as lipids or nucleic acids, is altered, and this then inform on whether the uptake of the EVs by receptor cells may be altered. This is particularly relevant in the context of neurodegenerative diseases, as these vesicles may not only play a role in the transmission of proteins but also in signaling cellular alterations taking place during the disease process.

Microglia and astrocytes are two important cell types in the brain that mediate neuroinflammatory processes and, thereby, playing an important role in the pathogenesis of neurodegenerative diseases (Kwon and Koh, 2020, Palpagama et al., 2019). Our study shows that the uptake ectosomes and exosomes can be taken up by microglial and astrocytic cells, and that both EV types elicited an increase in pro-inflammatory cytokines (IL-6, IL-1β and TNFα). Microglial cells adopted an activated phenotype and exhibited LC3 puncta in the cytoplasm and increase in p62 levels, indicating autophagy activation, possible for the clearance of the EVs. Astrocytes also displayed accumulation of EVs in the cytoplasm, with the EGFP signal being surrounded by LC3 staining. Interestingly, microglia cells were more sensitive to EV-treatment: while astrocytes tolerated 20*μ*g/mL, microglia tolerated only up to 10*μ*g/mL of EV protein. Ectosomes and exosomes containing aSyn-EGFP, EGFP-Tau or 25QHtt-EGFP were taken up at similar levels in microglia, and these were higher than those observed with 103QHtt-EGFP. Indeed, both wild-type and mutant Htt can influence vesicle transport in the secretory and endocytic pathways through associations with clathrin-coated vesicles (Velier et al., 1998). In contrast, astrocytes seemed to, in general, internalize more ectosomes, or degrade exosomes faster. Together, our findings demonstrate that EVs can be targeted by different types of glial cells, and that their uptake and effects are likely correlated with the EV type and content. These findings provide new insights into molecular mechanisms of intercellular communication.

We also observed that ectosomes and exosomes are taken up by primary cortical neurons, as previously described (Bras et al., 2021). The internalization ratio was similar for EVs with aSyn, Tau or Htt, but was considerably lower than that observed with microglia and astrocytes. The potential effects of EVs on neuronal network activity are still unclear. In this context, we demonstrate that spontaneous neuronal function can be modulated by ectosomes and exosomes, and that EV internalization is associated with a disruption of the typical synchronized bursting activity, resulting mostly in lower and less organized spiking activity. Interestingly, these alterations are mainly correlated with the EV subtype, and not with the presence of aSyn, Tau or Htt, although slight differences could be perceived between 25QHtt-EGFP and 103QHtt-EGFP (Bras et al., 2021).

We also demonstrate that aSyn, Tau or Htt present in the cell media can modify the spontaneous activity in cortical neurons. The use of identical concentrations of monomeric protein allowed us to compare the consequences of aSyn, Tau and Htt internalization, at sub-cytotoxic concentrations. aSyn function has been associated with synaptic activity through the regulation of the vesicle pool (Volpicelli-Daley et al., 2011, Maroteaux et al., 1988, Fortin et al., 2004, Busch et al., 2014). Interestingly, incubation of cortical neurons with monomeric aSyn resulted in a reduction of the firing rate and in an increase in burst duration, without changing the coordinated network activity. These effects contrast with those reported with higher concentrations of extracellular aSyn or with aggregated assemblies that strongly reduce neuronal activity by disrupting synaptic transmission, thereby contributing to neuronal death (Hassink et al., 2018, Shrivastava et al., 2020). Remarkably, treatment of neurons with monomeric Tau results in increased neuronal activity and in robust and synchronized bursting activity, suggesting that neurons fire in a more regular and synchronized manner. Consistently, full-length monomeric Tau was previously described to be rapidly and efficiently internalized in healthy neurons, implying this might be part of a physiological, and not pathological, process (Evans et al., 2018). Treatment of neurons with 23QHtt or 43QHtt resulted in longer burst intervals and a reduction in the synchronized bursting activity. As expected, internalization of 43QHtt resulted in greater impairment in the coordinated network activity, correlated with the toxic effects of the polyglutamine expansion (Ren et al., 2009, Masnata et al., 2019, Yang et al., 2002). A detailed understanding of the mechanisms of internalization of monomeric and aggregated forms of aSyn, Tau and Htt will be invaluable for the development of potential therapies for preventing the interneuronal transfer of proteins, without interfering with the physiological transfer of non-pathogenic forms.

Overall, our systematic study compares the transfer of disease-related proteins through various cellular mechanisms, and between different cell types. In particular, we emphasize that protein release, either in a free form or in EVs, induces diverse effects in neighboring receptor cells, and that great care is important when considering the development of therapeutic strategies to avoid interfering with normal physiological intercellular communication.

## Acknowledgments

We thank Prof. Dr. Ray Truant (McMaster University, Ontario, Canada) for kindly providing the Htt antibody used in the immunoblot experiments [anti-N17 (1-8)]. We also thank Prof. Dr. Hilal Lashuel (EPFL, Switzerlenad), for kindly providing the recombinant 23QHtt and 43QHtt, and to Prof. Dr. Flaviano Giorgini (Leicester University, Leicester, United Kingdom) for providing the Htt lentiviral vectors. We thank Sabine König and Uwe Plessmann from Max Planck Institute for Biophysical Chemistry (Göttingen, Germany) and Christof Lenz from the Core Facility Proteomics at University Medical Center Göttingen (Göttingen, Germany) for assisting with mass spectrometry analysis. We thank Dr. Nicolás Lemus (Göttingen, Germany) for support with flow cytometry experiments.

## Funding

TFO is supported by European Union’s Horizon 2020 research and innovation program under grant agreement No. 721802 (SynDegen), and by the Deutsche Forschungsgemeinschaft (DFG, German Research Foundation) under Germany’s Excellence Strategy - EXC 2067/1-390729940, by SFB1286 (B8). TG is supported by the European Research Council (ERC) under the European Union’s Horizon 2020 research and innovation programme (grant agreement number 724822). This work was partly supported by the Göttingen Graduate School for Neurosciences, Biophysics, and Molecular Biosciences (DFG grant GSC 226/4).

## Author contributions

T.F.O. and I.C.B. conceived the study. I.C.B performed all the cell culture, molecular biology, and imaging experiments. M.H.K and I.C.B established the spike sorting framework, performed MEA experiments and data analysis. E.V prepared the microglia samples for qPCR, imaging and the recombinant aSyn. W.M performed the immuno-EM experiments in the stable cell lines. D.R performed the EM experiments in EVs. E.G performed the cloning and prepared the lentiviral vectors. C.F. prepared the lentiviral vectors. I.P and H.U performed the mass spectrometry experiments. M.Z. provided the recombinant full-length 2N4R Tau. T.G. provided methodology and resources for the MEA experiments. I.C.B analyzed and interpreted the data. I.C.B generated the graphs and figures. I.C.B. and T.F.O. wrote the manuscript. T.F.O. supervised the work.

## Declaration of Interests

The authors declare no competing interests.

## Supplementary figure legends

**Supplementary Figure 1. Different disease-related proteins are secreted to the extracellular space. (A)** Representative images of HEK cells transfected with plasmids encoding untagged 22QHtt, 72QHtt, aSyn or Tau. Control cells were transfected with an empty plasmid. Scale bar 5 µm. **(B)** Western blots showing the protein levels in the lysates and released to the cell media of the different cells. Quantifications were normalized to total protein levels using MemCode. **(C-E)** LDH measurements confirm the absence of cell toxicity and cell death in **(C)** cells transfected with disease-relate proteins without tag, **(D)** in HEK cells stably expressing 25QHtt-EGFP, 103QHtt-EGFP, aSyn-EGFP or EGFP-Tau, and **(E)** in primary cortical neurons expressing 25QHtt-EGFP, 103QHtt-EGFP, aSyn-EGFP or EGFP-Tau. Data from at least three independent experiments for each condition. Data from at least three independent experiments for each condition. Significant differences were assessed by one-way ANOVA followed by multiple comparisons with significance between groups corrected by Bonferroni procedure.

**Supplementary Figure 2. Purification and characterization of secreted EVs using differential centrifugation. (A)** MemCode staining demonstrates the total protein levels present in each fraction. Exosome-depleted cell media was collected from HEK cells after 24 hours and subsequently centrifuged at different speed. **(B)** Whole-mount electron microscopy analysis of each pellet showing representative images of ectosomes and exosomes (scale bar 100 nm). **(C)** Nanoparticle tracking analysis (NTA) measurements of particle concentrations and average size distributions of ectosomes and exosomes. Average is represented with the filled line while each dotted line represents one biological replicate. Yellow dots represent exosomes measurements, while orange dots represent ectosomes measurements. Data from at least three independent experiments for each condition. **(D)** Proteomic analyses of ectosomes and exosomes using label-free quantitative mass spectrometry demonstrates the enrichment of specific protein markers in each fraction. Yellow dots represent the proteins enrichment in exosomes, while orange dots represent enrichment in ectosomes. Dots above the volcano plot line represent proteins for which differences were significant (false discovery rate [FDR] <0.1). Data represented in “t-test Difference (Ectosomes - Exosomes)” vs. “-Log t-test p-value” from 5 independent samples for each group. Data analyses were performed using Perseus software.

**Supplementary Figure 3. Disease-related proteins are enriched in ectosomes. (A)** Immunoblots of ectosomes and exosomes purified from the media of HEK cells stably expressing 25QHtt-EGFP, 103QHtt-EGFP, aSyn-EGFP or EGFP-Tau for 24 hours. Equal quantities of protein were separated on SDS-PAGE gels, and membranes were blotted with the indicated antibodies. Protein levels were normalized to total protein levels using Memcode. Data from at least three independent experiments for each condition. Significant differences were assessed by two-tailed unpaired t test comparison and are expressed as mean ± SD, *p<0.05, **p<0.01. **(B-C)** Proteomic analyses of ectosomes and exosomes using label-free quantitative mass spectrometry demonstrates the enrichment of 25QHtt-EGFP, 103QHtt-EGFP, aSyn-EGFP and EGFP-Tau in ectosomes **(B)** and exosomes **(C)** compared with the control (proteins are identified in green). **(B)** Data represented in “t-test Difference (Ectosomes – Ectosomes 25QHtt-EGFP/ 103QHtt-EGFP/ aSyn-EGFP/ EGFP-Tau)” vs. “-Log t-test p-value” from 3 independent samples for each group. **(C)** Data represented in “t-test Difference (Exosomes – Exosomes 25QHtt-EGFP/ 103QHtt-EGFP/ aSyn-EGFP/ EGFP-Tau)” vs. “-Log t-test p-value” from 3 independent samples for each group. Dots above the volcano plot line represent proteins for which differences were significant (false discovery rate [FDR] <0.1). Data analyses were performed using Perseus software.

**Supplementary Figure 4. Ectosomes and exosomes containing disease-related proteins are internalized by microglial cells. (A)** Ectosomes and exosomes were applied to microglial cultures at a concentration of 10*μ*g/mL for 24 hours. Cells were immunostained for LC3 (red) and Iba1 (grey). Scale bar 10 µm. **(B)** Ectosomes and exosomes were labelled with Alexa Fluor 633 C5-maleimide (gray) and applied to microglial cultures at a concentration of 10*μ*g/mL for 24 hours. Cells were immunostained for Iba1 (red). Scale bar 10 µm. **(C)** Ectosomes and exosomes containing 25QHtt-EGFP, 103QHtt-EGFP, aSyn-EGFP or EGFP-Tau were labelled with Alexa Fluor 633 C5-maleimide (gray) and applied to microglial cultures at a concentration of 10*μ*g/mL for 24 hours. Cells were immunostained for Iba1 (red). Scale bar 10 µm. **(D)** EV internalization was evaluated through imaging analysis by measuring fluorescence intensity and cell area. Data from at least three independent experiments for each condition. Significant differences were assessed by one-way ANOVA followed by multiple comparisons with significance between groups corrected by Bonferroni procedure. Differences were considered to be significant for values of p<0.05 and are expressed as mean ± SD, *p<0.05, **p<0.01.

**Supplementary Figure 5. Ectosomes and exosomes containing disease-related proteins are internalized by microglial cells. (A)** Ectosomes and exosomes containing 25QHtt-EGFP, 103QHtt-EGFP, aSyn-EGFP or EGFP-Tau were applied to microglial cultures at a concentration of 10*μ*g/mL for 24 hours. Immunoblot and protein quantifications of iNOS, Iba1 and APG5L/ ATG5. **(B)** EV treatment leads to the activation of the pro-inflammatory markers IL-6 and TNFα in microglia cells after 24hours. **(B)** LDH measurements confirm the absence of cell toxicity and cell death in the experiments. Data from at least three independent experiments for each condition. Significant differences were assessed by one-way ANOVA followed by multiple comparisons with significance between groups corrected by Bonferroni procedure. Differences were considered to be significant for values of p<0.05 and are expressed as mean ± SD, *p<0.05.

**Supplementary Figure 6. Ectosomes and exosomes containing disease-related proteins are internalized by astrocytic cells. (A)** Ectosomes and exosomes were applied to astrocytic cultures at a concentration of 20*μ*g/mL for 24 hours. Cells were immunostained for LC3 (red) and GFAP (grey). Scale bar 10 µm. **(B)** Ectosomes and exosomes were labelled with Alexa Fluor 633 C5-maleimide (gray) and applied to astrocytic cultures at a concentration of 20*μ*g/mL for 24 hours. Cells were immunostained for Iba1 (red). Scale bar 10 µm. **(C)** Ectosomes and exosomes containing 25QHtt-EGFP, 103QHtt-EGFP, aSyn-EGFP or EGFP-Tau were labelled with Alexa Fluor 633 C5-maleimide (gray) and applied to cell cultures at a concentration of 20*μ*g/mL for 24 hours. Cells were immunostained for Iba1 (red). Scale bar 10 µm. **(D)** EV internalization levels were evaluated by image analyses measuring fluorescence intensity and cell area. Data from at least three independent experiments for each condition. Significant differences were assessed by one-way ANOVA followed by multiple comparisons with significance between groups corrected by Bonferroni procedure. Differences were considered to be significant for values of p<0.05 and are expressed as mean ± SD, *p<0.05, **p<0.01, ***p<0.001, ****p<0.0001.

**Supplementary Figure 7. Ectosomes and exosomes containing disease-related proteins are internalized by astrocytic cells. (A)** Ectosomes and exosomes containing 25QHtt-EGFP, 103QHtt-EGFP, aSyn-EGFP or EGFP-Tau were applied into astrocytic cultures at a concentration of 20*μ*g/mL for 24 hours. Immunoblot and protein quantifications of iNOS, APG5L/ ATG5, p62, GFAP and LC3. **(B)** EV treatment leads to the activation of the pro-inflammatory markers IL-6, IL-β and TNFα in astrocytic cells after 24hours. **(C)** LDH measurements confirm the absence of cell toxicity and cell death in the experiments. Data from at least three independent experiments for each condition. Significant differences were assessed by one-way ANOVA followed by multiple comparisons with significance between groups corrected by Bonferroni procedure. Differences were considered to be significant for values of p<0.05 and are expressed as mean ± SD, *p<0.05, **p<0.01, ***p<0.001, ****p<0.0001.

**Supplementary Figure 8. Ectosomes and exosomes containing disease-related proteins are internalized by primary cortical neurons. (A)** Ectosomes and exosomes were applied to primary cortical neurons 20*μ*g/mL for 24 hours. Cells were immunostained for LC3 (red) and GFAP (grey). Scale bar 5µm. **(B)** Ectosomes and exosomes were labelled with Alexa Fluor 633 C5-maleimide (gray) and applied to neuronal cultures at a concentration of 20*μ*g/mL for 24 hours. Cells were immunostained for MAP2 (red). Scale bar 5µm. **(C)** Ectosomes and exosomes containing 25QHtt-EGFP, 103QHtt-EGFP, aSyn-EGFP or EGFP-Tau were applied to primary cortical cultures at a concentration of 20*μ*g/mL for 24 hours. Immunoblot and protein quantifications of PSD95, synaptophysin, MAP2, APG5L/ ATG5, p62, and LC3. **(D)** LDH measurements confirm the absence of cell toxicity and cell death in the experiments. Data from at least three independent experiments for each condition. Significant differences were assessed by one-way ANOVA followed by multiple comparisons with significance between groups corrected by Bonferroni procedure. Differences were considered to be significant for values of p<0.05 and are expressed as mean ± SD.

## Materials and Methods

### Ethics statement

Mice (C57BL6/J#00245 background) were bred and maintained under specific pathogen free conditions in the animal facility of the University Medical Center Göttingen (Göttingen, Germany). All animal procedures were performed in accordance with the European Community (Directive 2010/63/EU), institutional and national guidelines (license numbers 19.3213 and T 20.7).

### Primary cultures

#### Neuronal cultures

C57BL6/J#00245 wild-type E15.5 embryos from the animal facility of the University Medical Center Göttingen (Göttingen, Germany) were used for the preparation of primary cortical neuronal cultures, as previously described (Szego et al., 2019). In detail, pregnant animals were anesthetized using carbon dioxide intoxication and the embryos extracted from the uterus. After removal of the meninges, cortex was dissected under a stereomicroscope and the tissue was transferred to ice-cold 1x Hanks’ balanced salt solution (CaCl_2_ and MgCl_2_ free) (HBSS; Gibco Invitrogen, CA, USA) supplemented with 0.5% sodium bicarbonate solution (Sigma-Aldrich, MO, USA). Trypsinization was performed at 37°C for 15 minutes (min) (100*μ*L of 0.25% trypsin; Gibco Invitrogen, CA, USA), and the reaction was stopped with the addition of 100*μ*L fetal bovine serum (FBS; Anprotec, Bruckberg, Germany) and 100*μ*L DNase I (0.5 mg/mL; Roche, Basel, Switzerland). After dissociation, the cell suspension was centrifuged at 300x*g* for 5 min and then resuspended in pre-warmed neurobasal medium (Gibco Invitrogen, CA, USA) supplemented with 2% B27 supplement (Gibco Invitrogen, CA, USA), 0.25% GlutaMAX (Gibco Invitrogen, CA, USA) and 1% penicillin-streptomycin (PAN Biotech, Aidenbach, Germany). Cells were seeded on coverslips coated with poly-L-ornithine (0.1 mg/mL in borate buffer) (PLO; Sigma-Aldrich, MO, USA) or culture plates (Corning, Merck, Darmstadt, Germany) for immunocytochemistry and western blot experiments. Cells were maintained at 37°C with 5% CO^2^, and fresh medium was added every 3-4 days.

#### Microglial cultures

Primary microglia were obtained from mixed glial cell cultures from C57BL6/J#00245 wild-type P0 pups from the animal facility of the University Medical Center Göttingen (Göttingen, Germany), as previously described (Garcia-Agudo et al., 2019). Briefly, meninges were removed from the isolated brains and the tissue was collected into ice cold 1x HBSS (Gibco Invitrogen, CA, USA). The supernatant was removed, the brains were washed 3 times with HBSS solution (without Ca^2+^, Mg^2+^ and phenol) (PAN Biotech, Aidenbach, Germany) and incubated with 0.05% trypsin-EDTA (PAN Biotech, Aidenbach, Germany) in a water bath at 37°C during 10 min. Trypsin was aspirated and the digestion was stopped by adding 0.5 mg/mL DNase I (Roche, Basel, Switzerland) in microglia medium [Dulbecco’s modified Eagle’s medium (DMEM; PAN Biotech, Aidenbach, Germany) supplemented with 0.5% penicillin-streptomycin (PAN Biotech, Aidenbach, Germany) and 10% FBS (Anprotec, Bruckberg, Germany)]. Tissue was shortly incubated for 2-3 min at 37°C in the water bath and carefully homogenized into single-cell suspensions with a glass pipette. Suspension was centrifuged for 10 min at 800x*g*, the supernatant was discarded, and the pellet was resuspended in microglia medium. Cell suspension was plated into T75 flask (Corning, Merck, Darmstadt, Germany) and incubated overnight at 37°C and 5% CO^2^. On the following day, 2 days *in vitro* (DIV2), the cells were washed 3 times with pre-warmed HBSS solution (PAN Biotech, Aidenbach, Germany), 1 time with microglia medium and incubated with new medium until the next day. On DIV3, cell medium was replaced once more. On DIV5, the culture was stimulated by substitution of one third of the medium with L929 medium previously prepared in aliquots (see description in the cell lines section). The first harvest was performed on DIV8 by mild shaking and collection of the floating microglial cells. Medium was then centrifuged for 10 min at 800x*g*, the pellet resuspended in microglia medium, and cells were plated into culture plates previously coated with PLO (0.1 mg/mL in borate buffer) (Sigma-Aldrich, MO, USA) for immunocytochemistry, western blot, and gene expression studies. The culture was re-stimulated with new medium containing one third of L929 medium and incubated during 2-3 days until the next harvest. Three harvests were performed for the same culture.

#### Astrocytic cultures

Primary astrocytic cultures were prepared from C57BL6/J#00245 wild-type cerebral cortices of P0 pups from the animal facility of the University Medical Center Göttingen (Göttingen, Germany), as previously described (Paiva et al., 2019). After decapitation, meninges were removed, and the cortex was isolated from the brains and kept in ice cold 1x HBSS (Gibco Invitrogen, CA, USA). Tissue was digested in a fresh prepared solution of 0.25% trypsin (Gibco Invitrogen, CA, USA), 0.5 mg/mL DNase I (Roche, Basel, Switzerland), 1mM EDTA (Carl Roth, Karlsruhe, Germany), 10mM HEPES (Gibco Invitrogen, CA, USA), 2mg/mL bovine serum albumin (BSA) (Sigma-Aldrich, MO, USA) in 1x HBSS (Gibco Invitrogen, CA, USA) at 37°C for 20 min. The reaction was stopped by adding astrocytes medium containing DMEM (PAN Biotech, Aidenbach, Germany) supplemented with 10% FBS (Anprotec, Bruckberg, Germany), 1% penicillin-streptomycin (PAN Biotech, Aidenbach, Germany) and 25mM HEPES (Gibco Invitrogen, CA, USA), followed by a short centrifugation at 800x*g* during 2 min. After aspiration of the supernatant, the tissue was dissociated with astrocytes medium until obtain a cell homogenate. After centrifugation at 800x*g* during 5 min, the pellet was resuspended in medium, and cells were plated in a T75 cm^2^ flask (Corning, Merck, Darmstadt, Germany). Mixed culture was incubated at 37°C with 5% CO^2^ for 3 days. At DIV1 cells were washed with warm HBSS solution (PAN Biotech, Aidenbach, Germany) and fresh media was added to the culture. The mixed astrocytic culture was agitated at DIV7 in an orbital plate shaker at 200 rpm for 40 min at 37°C. Cell culture agitation was repeated at DIV 9 for 2.5 hours (h) at 250 rpm (37°C). A third shake was performed on DIV10 at 350 rpm, overnight at 37°C. After each agitation step, the supernatant was aspirated and the remaining cells in the flask were washed 1 time with warm HBSS solution (PAN Biotech, Aidenbach, Germany) and fresh media was added to the cells. Cells were detached at DIV11 using 0.25% trypsin (Gibco Invitrogen, CA, USA) at 37°C for 5 min. Culture medium was added to stopped trypsin action, and the cell suspension was collected and centrifuged at 800x*g* for 5 min. Cell pellet was resuspended in medium and cells were seeded with fresh medium on plates or coverslips (Corning, Merck, Darmstadt, Germany) coated with PLO (0.1 mg/mL in borate buffer) (Sigma-Aldrich, MO, USA) for immunocytochemistry, western blot and gene expression studies at the appropriate density.

### Cell lines

#### Mouse fibroblast L929 cells

Generation of L929-conditioned medium as performed as previously described (Garcia-Agudo et al., 2019). L929 mouse fibroblast cells (kindly provided by Prof. Dr Hannelore Ehrenreich, MPI-EM, Göttingen, Germany) were plated into a T175cm^2^ cell culture flask (Corning, Merck, Darmstadt, Germany) with 100mL culture medium for 7 days containing DMEM (PAN Biotech, Aidenbach, Germany) supplemented with 10% FBS (Anprotec, Bruckberg, Germany) and 1% penicillin-streptomycin (PAN Biotech, Aidenbach, Germany) at 37°C with 5% CO^2^. The media was then collected, sterilized by filtration with 0.22µm filter (Sartorius, Göttingen, Germany) and stored at −20°C. The aliquots were freshly thawed when used for stimulation of the microglial cell culture.

#### Human embryonic kidney cells

Human embryonic kidney (HEK) 293 cells (ATTC, VA, USA) were maintained in DMEM medium (PAN Biotech, Aidenbach, Germany) supplemented with 10% FBS (Anprotec, Bruckberg, Germany) and 1% penicillin-streptomycin (PAN Biotech, Aidenbach, Germany) at 37°C in a 5% CO^2^ atmosphere.

### Transfection protocol

HEK 293 cells were seeded in cell culture plates (Corning, Merck, Darmstadt, Germany) in the day before at the appropriate density. Transfection protocol was performed using Metafectene (Biotex, TX, USA) according to the manufacturer’s instructions and using equimolar amounts of the plasmids of interest. The following plasmids used for the transfection protocol: pcDNA 3.1, pcDNA 3.1-aSyn, pcDNA 3.1-Tau (4R2N), pcDNA 3.1-22QHtt exon 1 (1-90, CAG, ID CHDI-90000027, Coriell Institute), pcDNA 3.1-72QHtt exon 1 (1-90, CAG, ID CHDI-90001882-1, Coriell Institute), mCitrine-USP19 (Plasmid #78593, Addgene), mCitrine-USP19 C506S (Plasmid #78594, Addgene). After 4h of transfection, fresh medium was added to the cells (as described above in the cell lines section). Cells were collected or stained for further western blot or immunocytochemistry analysis after 24h or 72h as described in each experiment section. USP19 plasmids were kindly provided by Prof. Dr. Yihong Ye through Addgene (Lee et al., 2016), and Htt plasmids were kindly provided by NIGMS Human Genetic Cell Repository at the Coriell Institute for Medical Research.

### Lentivirus production

Production of FUGW-aSyn-EGFP, FUGW-EGFP-Tau, pRRL-CMV-25QHtt-EGFP-PRE-SIN and pRRL-CMV-103QHtt-EGFP-PRE-SIN lentivirus was performed as previously described (Tiscornia et al., 2006). Htt lentiviral vectors were kindly provided by Prof. Dr. Flaviano Giorgini (Leicester University, Leicester, United Kingdom) (Kwan et al., 2012). HEK 293 cells were seeded in culture plates (Corning, Merck, Darmstadt, Germany) and incubated in DMEM (PAN Biotech, Aidenbach, Germany) supplemented with 10% FBS (Anprotec, Bruckberg, Germany) and 1% penicillin-streptomycin (PAN Biotech, Aidenbach, Germany) overnight at 37°C with 5% CO^2^. On the following day, cells were incubated with DMEM with 2% FBS (Anprotec, Bruckberg, Germany), transfection medium for 5 h before transfection. Cells were transfected using calcium phosphate (CaPO_4_) precipitation method with a plasmid mix [57.9*μ*g vesicular stomatitis virus glycoprotein (VSVG) packing virus, 144*μ*g of Delta 8.9 packaging virus, and 160*μ*g of the plasmid of interest]. The DNA mix was added to 6 mL of 1x BBS (50 mM BES, 280 mM NaCl, 1.5 mM Na_2_HPO_4_) and 0.36 mL CaCl_2_ (2.5M CaCl_2_) was added to the mixture in a vortex shaker in the dark under sterile conditions. Before adding the mixture to the cells, solution was incubated 20 min in the dark. On the following day, cells were incubated with Panserin (PAN Biotech, Aidenbach, Germany) supplemented with 1% of non-essential amino acids (MEM, Gibco Invitrogen, CA, USA) and 1% penicillin-streptomycin (PAN Biotech, Aidenbach, Germany). Viruses were harvested 72h after transfection and centrifuged at 3000x*g* for 15min at 4°C. The supernatant was filtrated through a 0.45µm filter (Sartorius, Göttingen, Germany) and incubated with 1x PEG solution (SBI System Bioscience, CA, USA) to pellet the viruses. Viruses were spin down by centrifugation at 1500x*g* during 30min (4°C) after 2 days of incubation at 4°C. The pellet was resuspended in 100*μ*L Panserin (PAN Biotech, Aidenbach, Germany).

### Lentiviral infections

For neuronal cortical cultures, cells were infected at DIV14 with FUGW-aSyn-EGFP, FUGW-EGFP-Tau, pRRL-CMV-25QHtt-EGFP-PRE-SIN and pRRL-CMV-103QHtt-EGFP-PRE-SIN and collected for further analyses at DIV19. Culturing conditions were the same as specified above (primary culture section).

Stable cell lines expressing alpha-synuclein (aSyn), Tau and huntingtin (Htt) were established by lentiviral infection of HEK 293 cells with FUGW-aSyn-EGFP, FUGW-EGFP-Tau, pRRL-CMV-25QHtt-EGFP-PRE-SIN and pRRL-CMV-103QHtt-EGFP-PRE-SIN. Cells were incubated during 5 days with the viruses and after 3 passages the infection rate was confirmed by microscopy (more than 90% of positive cells).

### Lactate dehydrogenase assay

Cytotoxicity in the cell cultures (primary cortical neurons and cell lines) was assessed using the cytotoxicity lactate dehydrogenase (LDH) detection kit according to the manufacturer’s instructions (Roche, Basel, Switzerland). Furthermore, culture medium was centrifuged at 500x*g* for 5 min to pellet cell debris before used in the experiments.

### Immunocytochemistry experiments

Primary cortical neurons and cell lines cultures after the treatments were washed with 1x PBS (PAN Biotech, Aidenbach, Germany) and fixed with 4% of paraformaldehyde solution (PFA) for 20 min (home-made) at room temperature (RT). PFA autofluorescence was quenched by incubation with 50 mM of ammonium chloride (NH_4_Cl) solution for 30 min at RT. Cells were permeabilized with 0.1% Triton X-100 (Sigma-Aldrich, MO, USA) for 10 min at RT, and then incubated with 2% BSA in PBS (NZYTech) blocking solution for 1h at RT. Incubation with primary antibody was performed overnight at 4°C [aSyn (1:1000, 610787, BD Biosciences), aSyn (phospho S129) (1:500, ab51253, Abcam), α-tubulin (1:1000, 302217, Synaptic Systems), Glial Fibrillary Acidic Protein (GFAP) (1:1000, Z0334, Dako), Htt (1:500, MAB5374, Merck Millipore), Iba1 (1:1000, ab5076, Abcam), LC3 (1:500, PM036, MBL International), MAP2 (1:1000, ab11267, Abcam), Synaptophysin (1:500, 101002, Synaptic Systems), Tau (1:1000, MN1000, Thermo Fisher Scientific)]. Subsequently, the cells were washed with 1x PBS (PAN Biotech, Aidenbach, Germany) and then incubated with fluorescence conjugated secondary antibodies for 2h at RT [Alexa Fluor 488 donkey (1:1000, A21206, A21202, A11055, A21208, Invitrogen), Alexa Fluor 555 donkey (1:1000, A31572, A31570, A21432, Invitrogen), Alexa Fluor 633 goat (1:1000, A21050, Invitrogen), Alexa Fluor 633 donkey (1:1000, A21082, Invitrogen) and Alexa Fluor 680 donkey (1:1000, A10043, Invitrogen)]. Lastly, nuclei were counter-stained with DAPI (Carl Roth, Karlsruhe, Germany) and mounted with mowiol (home-made) for imaging experiments.

### Western blots

Primary cortical neurons and cell lines cultures after the treatments were washed with 1x PBS (PAN Biotech, Aidenbach, Germany) and lysed in RIPA buffer [50mM Tris, pH 8.0, 0.15M NaCl, 0.1% SDS, 1.0% NP-40, 0.5% Na-Deoxycholate, 2mM EDTA, supplemented with protease and phosphatase inhibitors cocktail [(cOmplete^TM^ protease inhibitor and PhosSTOP^TM^ phosphatase inhibitor) (Roche, Basel, Switzerland)]. Lysate protein concentrations were determined using the Bradford protein assay (Bio-Rad, CA, USA). Samples containing 30µg of protein were denaturated for 5 min at 95°C, loaded into 12% SDS-PAGE gels and transferred to nitrocellulose membranes using iBlot 2 (Invitrogen, CA, USA). Membranes were incubated in blocking solution containing 5% skim milk (Sigma-Aldrich, MO, USA) in tris-buffered saline (pH 8) with 0.05% tween 20 (TBS-T). Primary antibody incubation was performed overnight at 4°C diluted in 5% BSA (Sigma-Aldrich, MO, USA) in TBS-T. The following primary antibodies were used in this study: Alix (1:1000, ab117600, Abcam), aSyn (1:3000, 610787, BD Biosciences), aSyn (phospho S129) (1:1000, ab51253, Abcam), α-tubulin (1:1000, 302217, Synaptic Systems), β-Actin (1:10000, A5441, Sigma-Aldrich), Annexin-A2 (1:1000, ab178677, Abcam), APG5L/ATG5 (1:1000, ab23722, Abcam), Flotilin-1 (1:1000, #18634, Cell Signaling), GFP (1:1000, 11814460001, Roche), Glial Fibrillary Acidic Protein (GFAP) (1:1000, Z0334, Dako), Huntingtin anti-N17 (1-8) [1:5000, kindly provided by Prof. Dr. Ray Truant (McMaster University, Ontario, Canada)], iNOS / NOS II (1:1000, 06-573, Upstate), Iba1 (1:1000, ab178846, Abcam), LC3 (1:1000, PM036, MBL International), MAP2 (1:1000, ab11267, Abcam), p62/SQSTM1 (1:1000, ab91526, Abcam), Synaptophysin (1:1000, 101002, Synaptic Systems), Tau (1:5000, A0024, Dako), USP19 (1:1000, A301-587A-M, Biomol). On the next day, the membranes were washed with TBS-T and incubated for 2 h with horseradish peroxidase (HRP) conjugated secondary antibodies [ECL™ Rabbit or Mouse IgG, HRP-linked whole antibody (1:10000, NA934V or NXA931, Amersham)]. Subsequently, membranes were washed with TBS-T and incubated with a chemiluminescent HRP substrate (Millipore, MA, USA) for bands visualization in a chemiluminescence system (Fusion FX Vilber Lourmat, Vilber, France). Intensities of specific bands were normalized to a protein loading control or to the total protein levels marked using MemCode™ Reversible Protein (Thermo Fisher Scientific, MA, USA).

### Native-PAGE

For native gel electrophoresis, samples were mixed with protein sample buffer (0.31M Tris HCl pH 6.8, 50% Glycerol, 0.4% Bromophenol blue). Samples were loaded on pre-cast vertical Serva gels 4–16% (Serva, Heidelberg, Germany) and ran according to the manufacturer’s instructions using the appropriate buffers [50 mM BisTris-HCl pH 7.0 for the anode buffer; 50 mM Tricin, 15 mM BisTris-HCl pH 7.0 for the cathode buffer] (Serva, Heidelberg, Germany). Proteins were transferred to nitrocellulose membranes using iBlot 2 (Invitrogen) and immunoblotting was performed as previously described in the western blot section.

### Isolation of extracellular vesicles

Isolation of ectosomes and exosomes was performed as previously described (Bras et al., 2021). HEK cells expressing the proteins of interest were grown in conditioned medium (depleted of FBS-derived exosomes). Briefly, DMEM (PAN Biotech, Aidenbach, Germany) supplemented with 20% FBS (Anprotec, Bruckberg, Germany) and 2% penicillin-streptomycin (PAN Biotech, Aidenbach, Germany) was centrifuged in polypropylene tubes (Optiseal; Beckman Coulter, CA, USA) in a fixed rotor (type 70 Ti, Beckman) during 16h at 100000x*g* (4**°**C), as previously described (Thery et al., 2006). The medium was subsequently diluted with DMEM medium (PAN Biotech, Aidenbach, Germany) for a final concentration of 10% FBS (Anprotec, Bruckberg, Germany) and 1% penicillin-streptomycin (PAN Biotech, Aidenbach, Germany). Cells were seeded in T75 cm^2^ flasks (Corning, Merck, Darmstadt, Germany) and incubated with fresh conditioned media during 24h at 80% confluency. The media was collected, and protease and phosphatase inhibitors [(cOmplete^TM^ protease inhibitor and PhosSTOP^TM^ phosphatase inhibitor) (Roche, Basel, Switzerland)] were added before centrifuging for 10 min at 300x*g* (4**°**C). A second centrifugation was performed at 2000x*g* for 20 min (4**°**C). Supernatant was transferred into ultra-clear tubes (Beckman Coulter, CA, USA) and ultracentrifuged in a swing rotor (TH-641 Sorvall; Thermo Fisher Scientific, MA, USA) during 90 min at 20000x*g* (4**°**C). Supernatant was carefully transferred into a new centrifuge tube and the pellet containing ectosomes was resuspended in ice cold PBS with protease and phosphatase inhibitors (PAN Biotech, Aidenbach, Germany). The medium was centrifuged in a swing rotor (TH-641 Sorvall; Thermo Fisher Scientific, MA, USA) during 90 min at 100000x*g* (4**°**C) to purify exosomes. The pellet was resuspended in ice cold PBS with protease and phosphatase inhibitors (PAN Biotech, Aidenbach, Germany). Ectosomal and exosomal pellets were diluted in PBS (PAN Biotech, Aidenbach, Germany) and centrifuged at the correspondent velocities to wash and concentrate the vesicles fractions. The pellets were resuspended in 100µl of ice cold 1x PBS with protease and phosphatase inhibitors (PAN Biotech, Aidenbach, Germany). Protein concentrations were determined by the BCA assay following the manufacturer’s instructions (Thermo Fisher Scientific, MA, USA).

### Labelling of extracellular vesicles

Extracellular vesicles were labelled with a fluorescent dye following a protocol previously described (Roberts-Dalton et al., 2017, Bras et al., 2021). Briefly, extracellular vesicles (60-100µg of total protein in 50*μ*L PBS) were incubated with Alexa Fluor 633 C5-maleimide dye (200*μ*g/mL; Invitrogen, Carlsbad, California, CA, USA) for 1h in the dark at RT. Samples were resuspended in 10mL PBS (PAN Biotech, Aidenbach, Germany) and centrifugated during 90 min at 20000x*g* (4**°**C) for ectosomes and 100000x*g* for 90 min for exosomes (4**°**C) in a swing rotor (TH-641 Sorvall; Thermo Fisher Scientific, MA, USA). As a control, PBS was similarly incubated with the dye to confirm the removal of the dye excess.

### NTA analysis

Extracellular vesicles size distribution and particle number were measured using nanoparticle tracking analysis (NTA) with NanoSight LM10 instrument (LM14 viewing unit equipped with a 532 nm laser) (NanoSight, Salisbury, UK). For each replicate, ectosomes and exosomes fractions were diluted in 1x PBS (PAN Biotech, Aidenbach, Germany) to a final volume of 400mL prior to analysis, according to the manufacturer’s recommendations (NanoSight, Salisbury, UK). Measurements were recorded using NTA 2.3 software with a detection threshold of 5, captured with a camera level of 16 at 25 °C, in videos of 5 x 60s repeated 4 times.

### Electron microscopy (EM)

Electron microscopy (EM) images from extracellular vesicles was performed following a protocol previously described (Bras et al., 2021). Isolated ectosomes and exosomes were bound to a glow discharged carbon foil covered grids. Samples were sained with 1% uranyl acetate *_(aq.)_* and then evaluated at RT using Talos L120C transmission electron microscope (Thermo Fisher Scientific, MA, USA).

### Immuno-EM

Immuno-EM images from HEK 293 cells expressing FUGW-aSyn-EGFP, FUGW-EGFP-Tau, pRRL-CMV-25QHtt-EGFP-PRE-SIN and pRRL-CMV-103QHtt-EGFP-PRE-SIN were performed following a protocol previously described (Slot and Geuze, 2007). Briefly, cells were fixed with 4% formaldehyde and 0.2% glutaraldehyde in 0.1M phosphate buffer. The cells were washed and then scraped from the dish in 0.1M phosphate buffer containing 1% gelatin, spun down, and resuspended in 10% gelatin in 0.1M phosphate buffer at 37°C. After spinning down again, the resulting pellets in gelatin were cooled on ice, removed from the tubes, and cut in small blocks. These blocks were infiltrated in 2.3M sucrose in 0.1M phosphate buffer, mounted onto aluminum pins for cryo-ultramicrotomy (Leica Microsystems, Vienna, Austria) and frozen in liquid nitrogen. Ultrathin cryosections were cut with a cryo-immuno diamond knife (Diatome, Biel, Switzerland) using a UC6 cryo-ultramicrotome (Leica Microsystems, Vienna, Austria) and picked up in a 1:1 mixture of 2% methylcellulose and 2.3M sucrose. For immuno-labeling, sections were incubated with 3H9 GFP primary antibody (#029762, ChromoTek, Planegg-Martinsried, Germany), followed by the secondary antibody (#112-4102, Rockland Immunochemicals, PA, USA) and then protein A-gold (10nm) (Cell Microscopy Core, Utrecht, Netherlands). Sections were analyzed with a LEO EM912 Omega (Zeiss, Oberkochen, Germany) and digital micrographs were obtained with an on-axis 2048×2048-CCD camera (TRS).

### Proteomic analyses of extracellular vesicles

For the proteomic analyses, ectosomes and exosomes samples were resuspended in Laemmli sample buffer and separated in SDS-PAGE gel, as previously described (Bras et al., 2021). Briefly, the entire lane was cut in 23 gel pieces and tryptically digested (Shevchenko et al., 2006). Extracted peptides were analyzed in technical replicates by liquid chromatography coupled to mass spectrometry (LC-MS) on a Dionex UltiMate 3000 RSLCnano system (Thermo Fisher Scientific, Waltham, USA) connected to an Orbitrap Fusion mass spectrometer (Thermo Fisher Scientific, Waltham, USA). A 43 min gradient ranging from 8% to 37% acetonitrile on an in-house packed C18 column was used to separate the peptides (75 µm x 30 cm, Reprosil-Pur 120C18-AQ, 1.9 µm, Dr. Maisch GmbH, Ammerbuch, Germany) at 300 nl/min flow rate. MS1 spectra were acquired with 120000 resolution (full width at half maximum, FWHM) and a scan range from 350 to 1600 m/z. Within a cycle time of 3s, precursor ions with a charge state between +2 and +7 were selected individually with a 1.6 m/z isolation window and were fragmented with a normalized collision energy of 35 by higher energy collisional dissociation (HCD). MS2 spectra were acquired in the ion trap with 20 % normalized AGC and dynamic injection time. Once selected precursors were excluded from another fragmentation event for 30s. Raw acquisition files were subjected to database search with Maxquant (version 1.6.2.10) (Cox and Mann, 2008) against the reference proteome of *Homo sapiens* (downloaded on 02/19/2019) and the GFP fusion proteins. Default settings were used unless stated differently below. Fractions were defined according to the cutting of the gel lanes and experiments were defined on the level of technical replicates. Unique and razor peptides were used for label-free quantification except for the GFP fusion constructs where all peptides were used due to the difficulty of peptide assignment and protein grouping related to the EGFP tag.

All the proteomic data analysis were performed with Perseus (version and 1.6.15.0) (Tyanova et al., 2016). The initial data was filtered for reverse hits, potential contaminants and hits only identified by site. Quantitative values were averaged across technical replicates ignoring missing values. A two-sample t-test was performed on biological replicates of the samples with an artificial within groups variance (s0) of 0.1 and a permutation-based FDR of 0.1 for multiple testing correction. These results were visualized in volcano plots.

### Secretion and internalization assays

The collection of the cell media and evaluation of proteins secretion was performed using an adapted protocol from a previous study (Trajkovic et al., 2017). Briefly, 1mL of medium from HEK cells expressing FUGW-aSyn-EGFP, FUGW-EGFP-Tau, pRRL-CMV-25QHtt-EGFP-PRE-SIN or pRRL-CMV-103QHtt-EGFP-PRE-SIN was collected after 24h and centrifuged for 5 min at 500x*g* at 4 °C to pellet cell debris. Supernatants were concentrated 10 times in an Amicon ultra 10K centrifugal filters (Millipore, MA, USA) following the manufacturer’s instructions. Only 30µL of the 100µL final volume was analyzed by western blot.

For the neuronal cell media, the DotBlot system was used to exclude the possibility that the proteins of interest could be retained in the centrifugal filters and change the results observed in the experiments. Briefly, 1mL of the neuronal cell media was collected from primary cortical neurons expressing FUGW-aSyn-EGFP, FUGW-EGFP-Tau, pRRL-CMV-25QHtt-EGFP-PRE-SIN or pRRL-CMV-103QHtt-EGFP-PRE-SIN and centrifuged for 5 min at 500x*g* (4°C). Only 300µL of the 1mL final volume was directly applied into the system in a nitrocellulose membrane (Bio-Rad, CA, USA).

For the internalization experiments, 1mL of medium from HEK cells expressing FUGW-aSyn-EGFP, FUGW-EGFP-Tau, pRRL-CMV-25QHtt-EGFP-PRE-SIN or pRRL-CMV-103QHtt-EGFP-PRE-SIN was collected after 24h and centrifuged for 5 min at 500x*g* (4°C). Afterwards, the media was added to naïve HEK cells for 72h.

### Flow cytometry experiments

As described in the previous section, naïve HEK cells were incubated during 72h with 1mL of cell media from HEK cells expressing FUGW-aSyn-EGFP, FUGW-EGFP-Tau, pRRL-CMV-25QHtt-EGFP-PRE-SIN or pRRL-CMV-103QHtt-EGFP-PRE-SIN. Subsequently, HEK cells were washed with 1x PBS and trypsinized (PAN Biotech, Aidenbach, Germany). Cell suspension was centrifuged during 5 min at 300x*g* (4°C). Cell pellet was resuspended in 1x PBS (PAN Biotech, Aidenbach, Germany) to remove residual cell media and centrifuged for 5 min at 300x*g* (4°C). The supernatant was removed by aspiration and the cell pellet was resuspended in 1 mL of ice cold 1x PBS (PAN Biotech, Aidenbach, Germany) with 0.1% of propidium iodide (Sigma-Aldrich, MO, USA). Cells without EGFP expression, treated with 0.1% triton-X (Sigma-Aldrich, MO, USA) or PBS alone were used as negative controls. 10 000 events were acquired on a FACSAria II flow cytometer (BD Biosciences, NJ, USA). Flow cytometric data were analyzed with FlowJo Analysis Software (BD Biosciences, NJ, USA).

### Treatment of cells with extracellular vesicles

Primary cortical neurons and astrocytes were treated with 20*μ*g/mL of ectosomes or exosomes resuspended in 1x PBS (PAN Biotech, Aidenbach, Germany) with protease and phosphatase inhibitors [(cOmplete^TM^ protease inhibitor and PhosSTOP^TM^ phosphatase inhibitor) (Roche, Basel, Switzerland)]. Due to high toxicity, microglial cells were treated only with 10*μ*g/mL of extracellular vesicles. Treatment was performed in cortical neurons at DIV14, and cells were fixed or collected for further analyses at DIV15 (24h treatment). Microglial and astrocytic cell cultures were treated in the day after their plating for 24h, and then fixed or collected for further analyses. Pro-inflammatory stimulation was evaluated by the exposition to lipopolysaccharide (LPS; Thermo Fisher Scientific, MA, USA). Microglia cells were treated with 50ng/mL and astrocytes with 100ng/mL of LPS (Thermo Fisher Scientific, MA, USA). Culturing conditions were the same as specified above in the primary culture section.

### Gene expression studies - RNA isolation and Quantitative real-time PCR

Total RNA was extracted from microglial and astrocytic cultures 24h after treatment using TRIzol Reagent according to the manufacturer’s instructions (Invitrogen, CA, USA). Reverse transcription of RNA to produce cDNA was performed using QuantiTect Reverse Transcription kit (Qiagen, MD, USA) following the protocol provided by the manufacturers. Quantitative real-time PCR (qPCR) analysis was performed on an Applied Biosystems Real-Time PCR Systems using SYBR Green Master Mix (Qiagen, MD, USA). The thermal cycler conditions were as following: 95°C for 10 min, then 40 cycles at 95°C for 15 s and 60°C for 25 s. Primers used in the quantitative real-time PCR experiments to evaluate the inflammatory markers were (mouse, sequence 5’ to 3’): IL-6 Forward-ATCCAGTTGCCTTCTTGGGACTGA, IL-6 Reverse-TAAGCCTCCGACTTGTGAAGTGGT, IL-10 Forward-GGTTGCCAAGCCTTATCGGA, IL-10 Reverse-CACTCTTCACCTGCTCCACT, IL-1β Forward-TCATTGTGGCTGTGGAGAAG, IL-1β Reverse-AGGCCACAGGTATTTTGTCG, TNFα Forward-CCCTCTCATCAGTTCTATGG, TNFα Reverse-GGAGTAGACAAGGTACAACC, β-actin Forward-GCGAGAAGATGACCCAGATC and β-actin Reverse-CCAGTGGTACGGCCAGAGG. Fold change expressions were calculated using the 2^−ΔΔCT^ method, with β-actin as a reference gene (Livak and Schmittgen, 2001). Quantification in the graphs show the normalized relative quantity (NRQ) values compared with the control.

### Treatment with recombinant monomeric protein

Primary cortical neurons were treated at DIV14 with recombinant monomeric protein until DIV19 (5 days). Final concentration of protein in the cultures was 100nM and PBS was employed as negative control. Recombinant aSyn, full-length 2N4R Tau, 23QHtt and 43QHtt were prepared as previously described (Dominguez-Meijide et al., 2020, Ukmar-Godec et al., 2020, Reif et al., 2018). Culturing conditions were the same as specified above (primary culture section).

### Multielectrode array experiments

Multi-electrode array (MEA) experiments were performed following standard protocols, as previously described (Khani and Gollisch, 2021, Bras et al., 2021). Primary cortical neuronal cultures were plated directly on 60MEA200/30iR-Ti-gr planar arrays (60 electrodes, 30µm electrode diameter, 200µm electrode spacing) (MultiChannel Systems, Reutlingen, Germany). The arrays were coated with poly-L-lysine overnight at 4°C (500µg/mL in borate buffer; PLL) (Sigma-Aldrich, MO, USA) and the next day with laminin for 1h at RT (5µg/mL in distilled water) (Sigma-Aldrich, MO, USA). Neuronal cells were plated directly on top of the electrodes and treated with EVs or recombinant proteins. For the EVs recordings, cells were incubated with 20µg/mL of ectosomes or exosomes containing the different disease-related proteins at DIV14 and recorded at DIV15, 24h after the treatment. In the recombinant monomeric protein experiments, cells were incubated with 100nM of aSyn, full Tau, 23QHtt exon 1 or 43QHtt exon 1 at DIV14 and recorded at DIV19, 5 days after the treatment. The neuronal activity was recorded using the MultiChannel MEA2100 system (MultiChannel Systems, Reutlingen, Germany) with temperature maintained at 35-37°C. Recordings started 10 min after translocation of the arrays from the incubator to the recording stage to avoid movement-induced artifacts, and the spontaneous activity was recorded for 5-10 min. The electrode signals were amplified, band-pass filtered (200Hz to 3kHz) and recorded digitally at 25kHz, using the MultiChannel Experimenter software (version 2.17.7.0) (MultiChannel Systems, Reutlingen, Germany). Spike sorting was carried out using a modified version of the Kilosort automatic sorting software, as previously described (Pachitariu et al., 2016b, Pachitariu et al., 2016a, Bras et al., 2021) (available at: https://github.com/MouseLand/Kilosort and https://github.com/dimokaramanlis/KiloSortMEA). Kilosort output was visually inspected and manually curated with the Phy2 software (https://github.com/cortex-lab/phy). Only spike clusters (“units”) with a well-separated spike waveform and a clear refractory period were included in the data analysis and considered as originated from individual neuronal cells. The spike clusters were pre-processed and analyzed using custom-made MATLAB scripts (Version: 9.7.0, R2019b; Mathworks, MA, USA). The raster plots, voltage traces, average firing rate and spike amplitude were measured from the spontaneous activity of each recorded cell. In all the recordings, we observed the occurrence of frequent bursts [groups of spikes occurring rapidly and consecutively with short inter-spike intervals, less than few tens of milliseconds (ms)], followed by quiescent periods longer than normal inter-spike intervals [generally several seconds (s) in our recordings]. The burst activity usually occurred synchronously for multiple cells over the array and in our analysis, and we focused on this population-wide synchronized bursts for further data analysis. To detect the concurrent bursts, the population firing rate was computed as a histogram (100ms bin size) of array-wide spiking activity. The peaks of the firing rate histogram were used to detect synchronous, array-wide bursts with at least 500 ms distance between two consecutive peaks. The peaks that were smaller than 1/5 of the largest peak were excluded as they do not correspond to array-wide synchronous activity. A time window of 650 ms around each peak (150 ms before to 500 ms after) was defined as the burst window (onset and offset of each burst). For each recorded cell, the spikes belonging to bursts were measured during the defined burst windows, and cells with fewer than six spikes across all their detected bursts were excluded from this analysis. From the detected bursts, the following parameters were calculated: (1) burst rate of the culture as the number of bursts per time over the duration of each recording; (2) inter-burst-interval as the time between the measured offset of a burst and the onset of the following burst, calculated for each pair of successive bursts in a recording; (3) burst duration for each cell as the time between the cell’s first and last spike during the burst window; (4) intra-burst-frequency as the rate of spikes occurring within a burst, averaged over all the detected bursts for each cell; and (5) percentage of spikes in bursts as the ratio of spikes occurring during bursts relative to the total number of spikes for each cell.

### Confocal microscopy imaging

Imaging was performed on a Leica SP5 confocal laser scanning microscope equipped with hybrid detectors using Application Suite X software with 100x immersion objective lenses (Leica Biosystems, Wetzlar, Germany). Samples were excited using 405 Diode, argon and helium–neon 633 lasers, pinhole = 1, 0.2 µm thickness Z stacks and 2 averaging line-by-line. The acquisition settings were optimized to avoid underexposure and oversaturation effects and kept equal throughout image acquisition of the samples.

### Quantifications and Statistical Analyses

Analyses of the images was performed using ImageJ software (National Institutes of Health) (Schindelin et al., 2012). The EVs uptake in neuronal cells was measured by the ration of the EVs signal area and cell area from different isolated areas chosen randomly within regions containing EVs signal. All data are presented as mean ± SD. Data from at least three independent experiments and each replicate represents one independent experiment. To assess differences between two groups, two-tailed unpaired student t-test was performed using GraphPad Prism 9 software (GraphPad, CA, USA). To assess differences between more than two groups, significant differences were assessed by one-way ANOVA followed by multiple comparisons with significance between groups corrected by Bonferroni procedure using GraphPad Prism 9 software (GraphPad, CA, USA). Differences were considered to be significant for values of p<0.05 and are expressed as mean ± SD. For mass spectrometry, the spectral count differences between samples were considered to be significant for FDR values<0.1 (see proteomic analyses section).

## Notes

### Competing Interest Statement

The authors have declared no competing interest.

