## Supplementary figures for "Common molecular mechanisms underlie the transfer of alpha-synuclein, Tau and huntingtin and modulate spontaneous activity in neuronal cells"

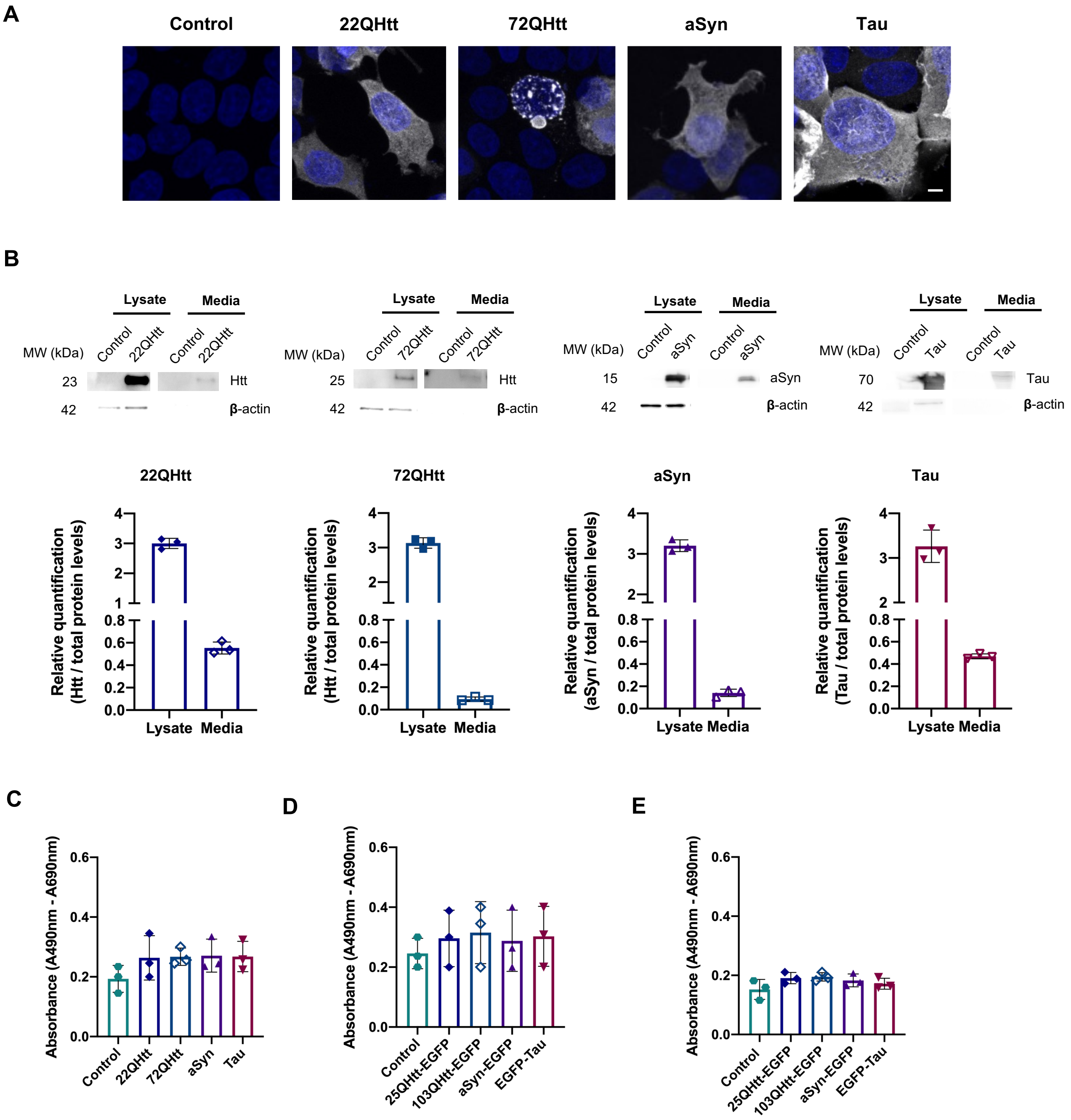


**Supplementary Figure 1. Different disease-related proteins are secreted to the extracellular space.**

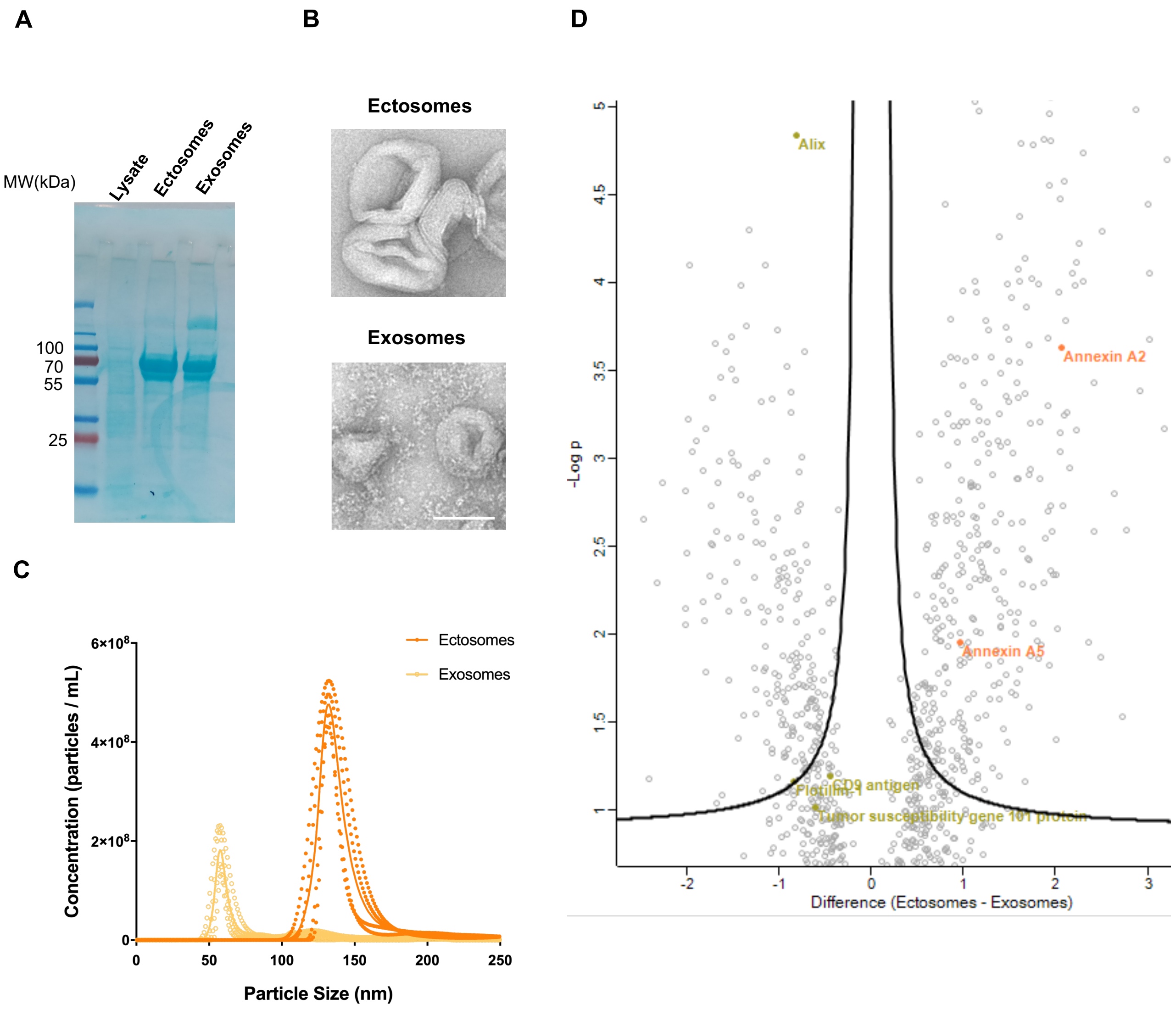


**Supplementary Figure 2. Purification and characterization of secreted EVs using differential centrifugation.**

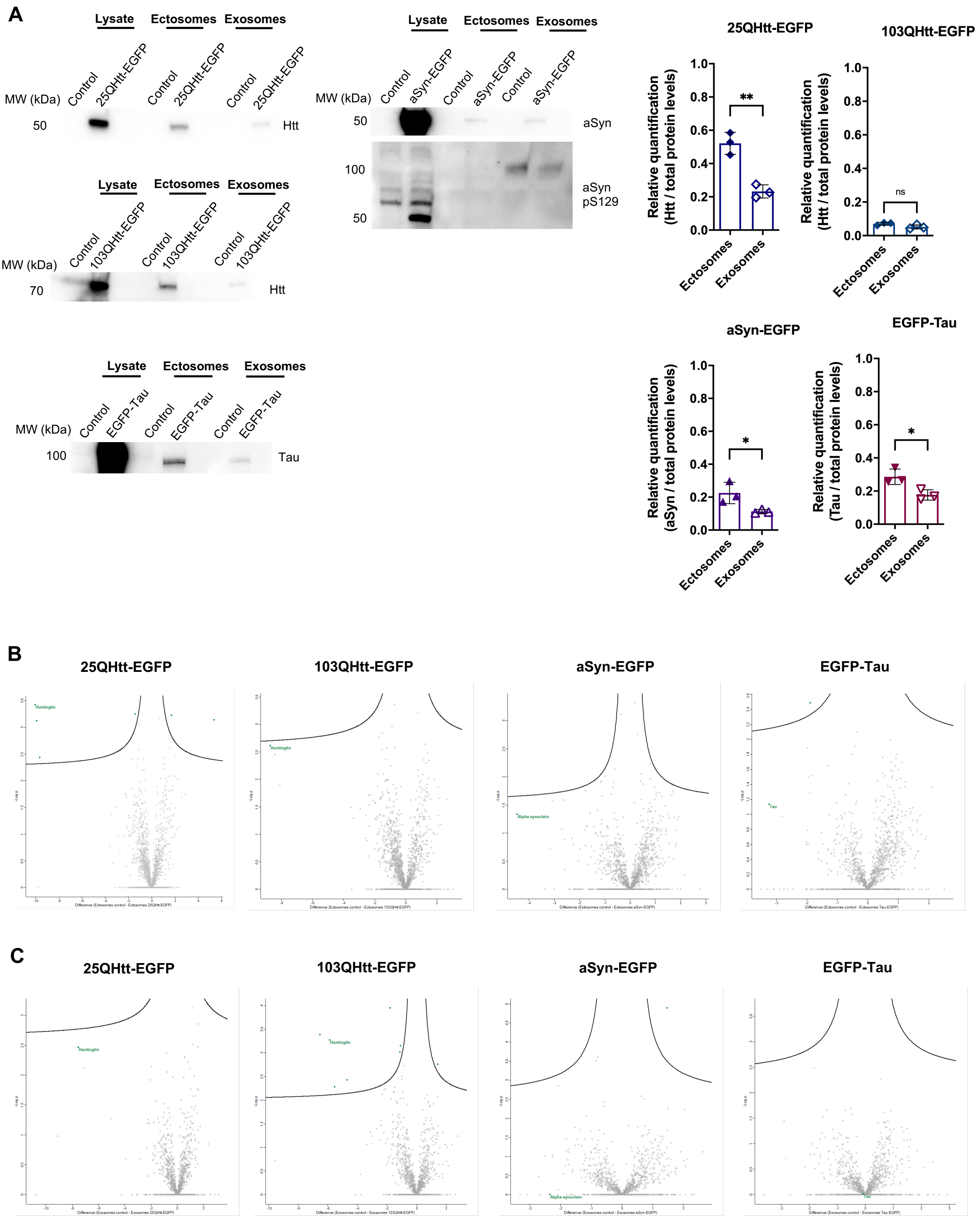


**Supplementary Figure 3. Disease-related proteins are more enriched in ectosomes.**

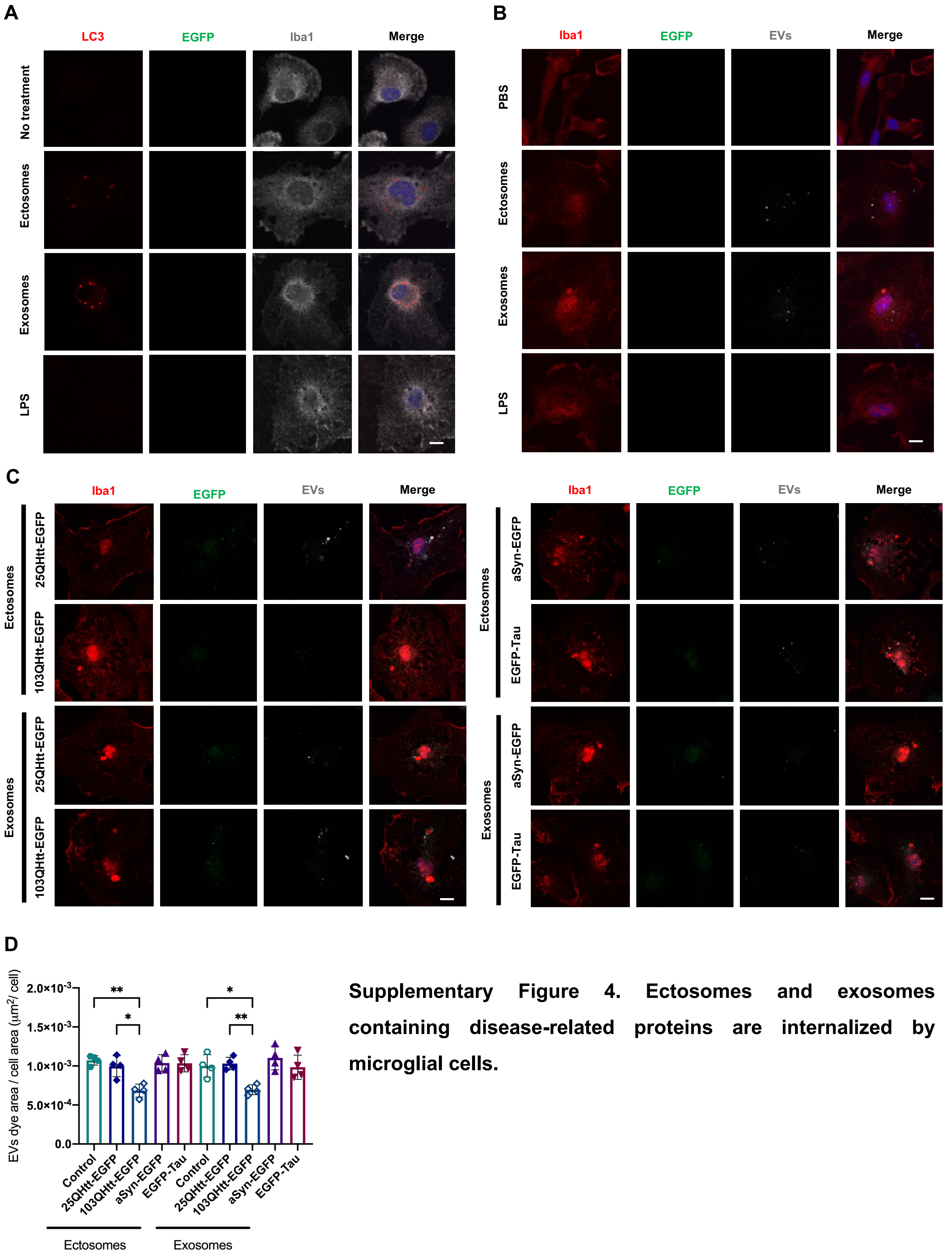


**Supplementary Figure 4. Ectosomes and exosomes containing disease-related proteins are internalized by microglial cells.**

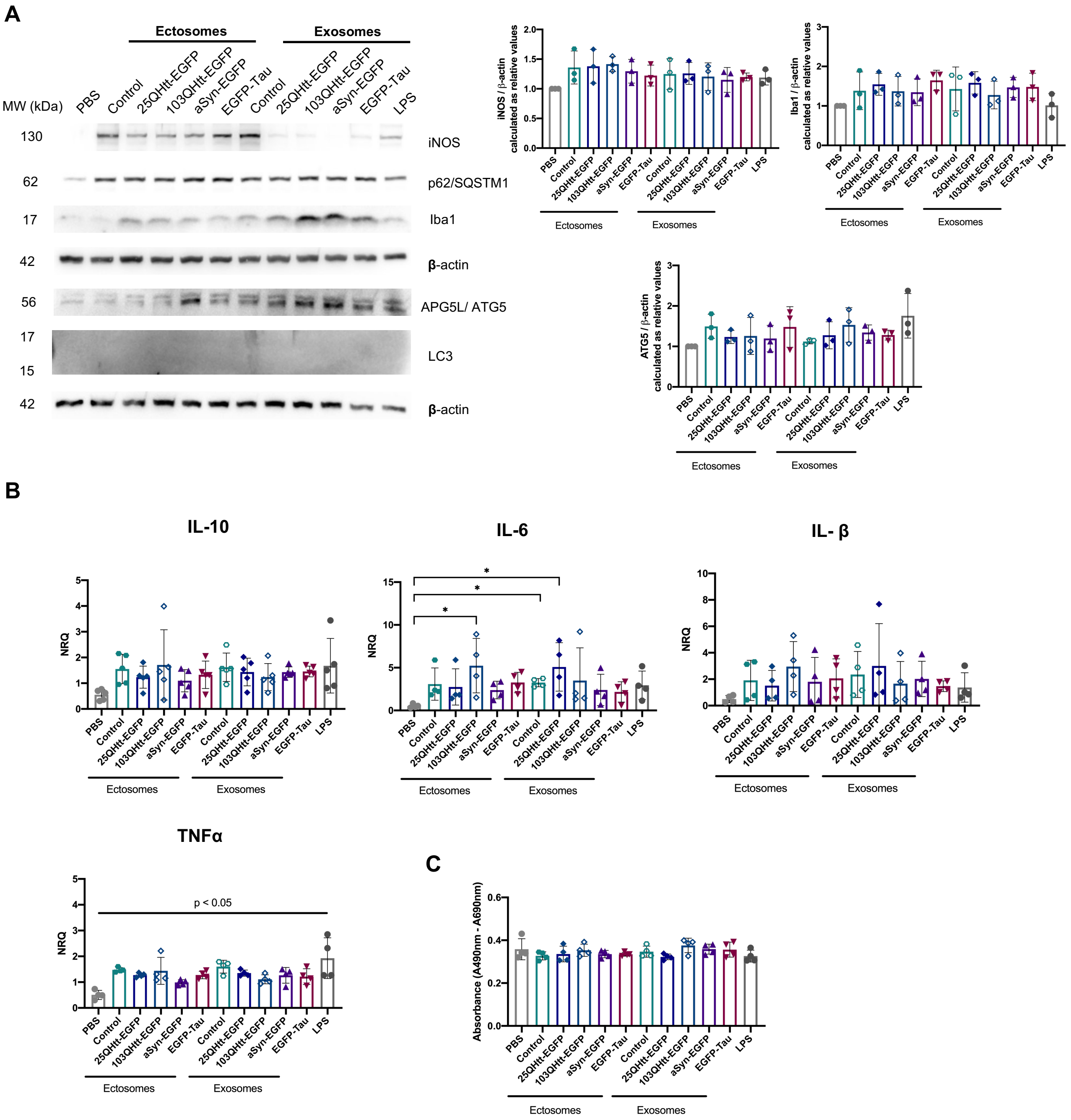


**Supplementary Figure 5. Ectosomes and exosomes containing disease-related proteins are internalized by microglial cells.**

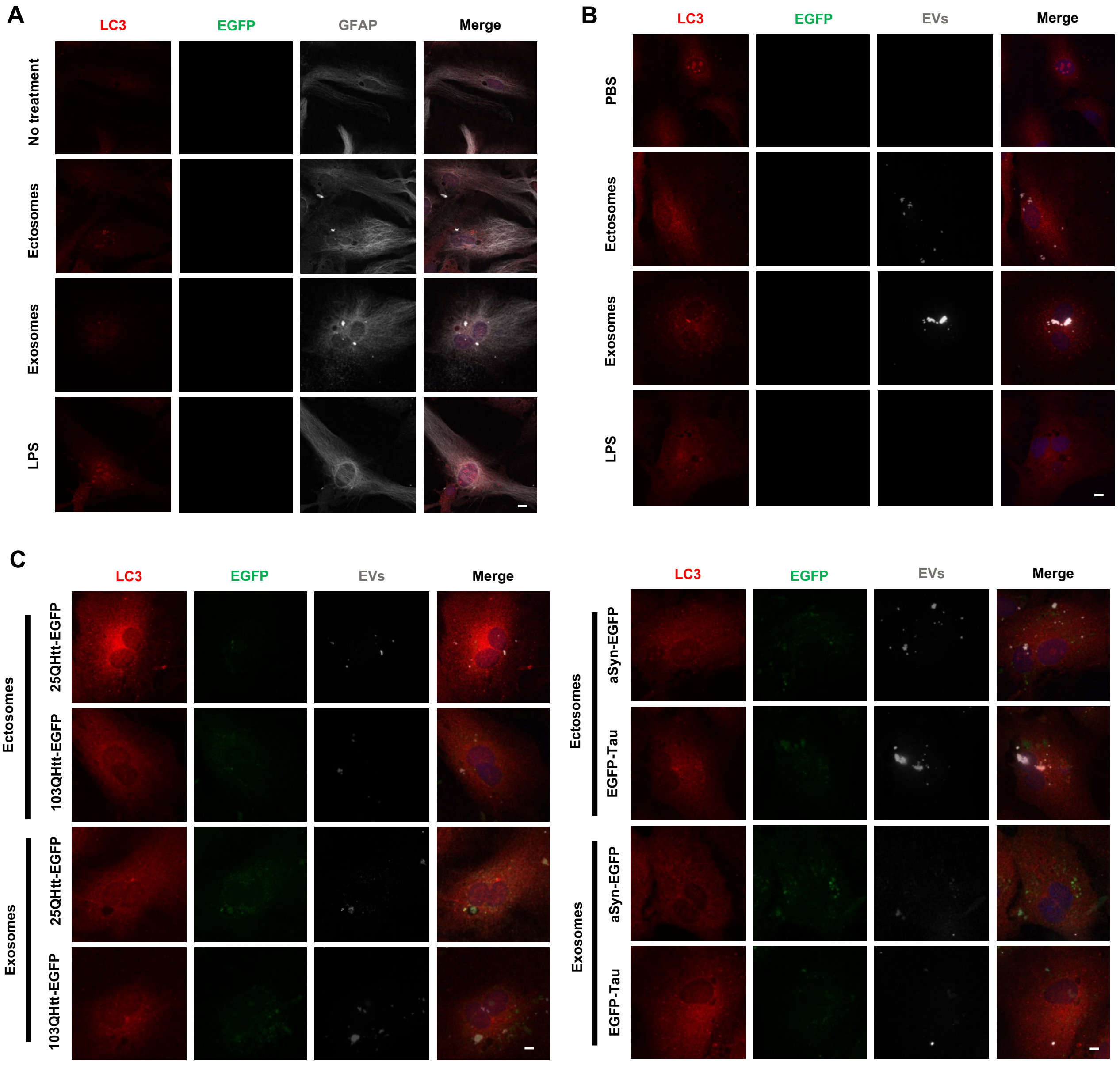


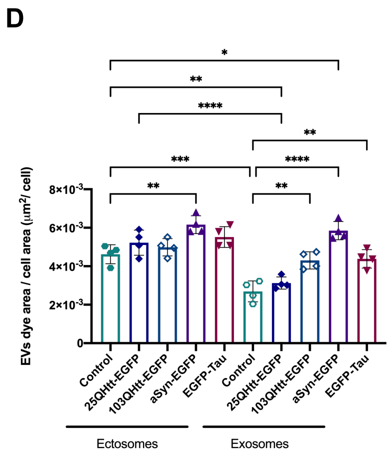


**Supplementary Figure 6. Ectosomes and exosomes containing disease-related proteins are internalized by astrocytic cells.**

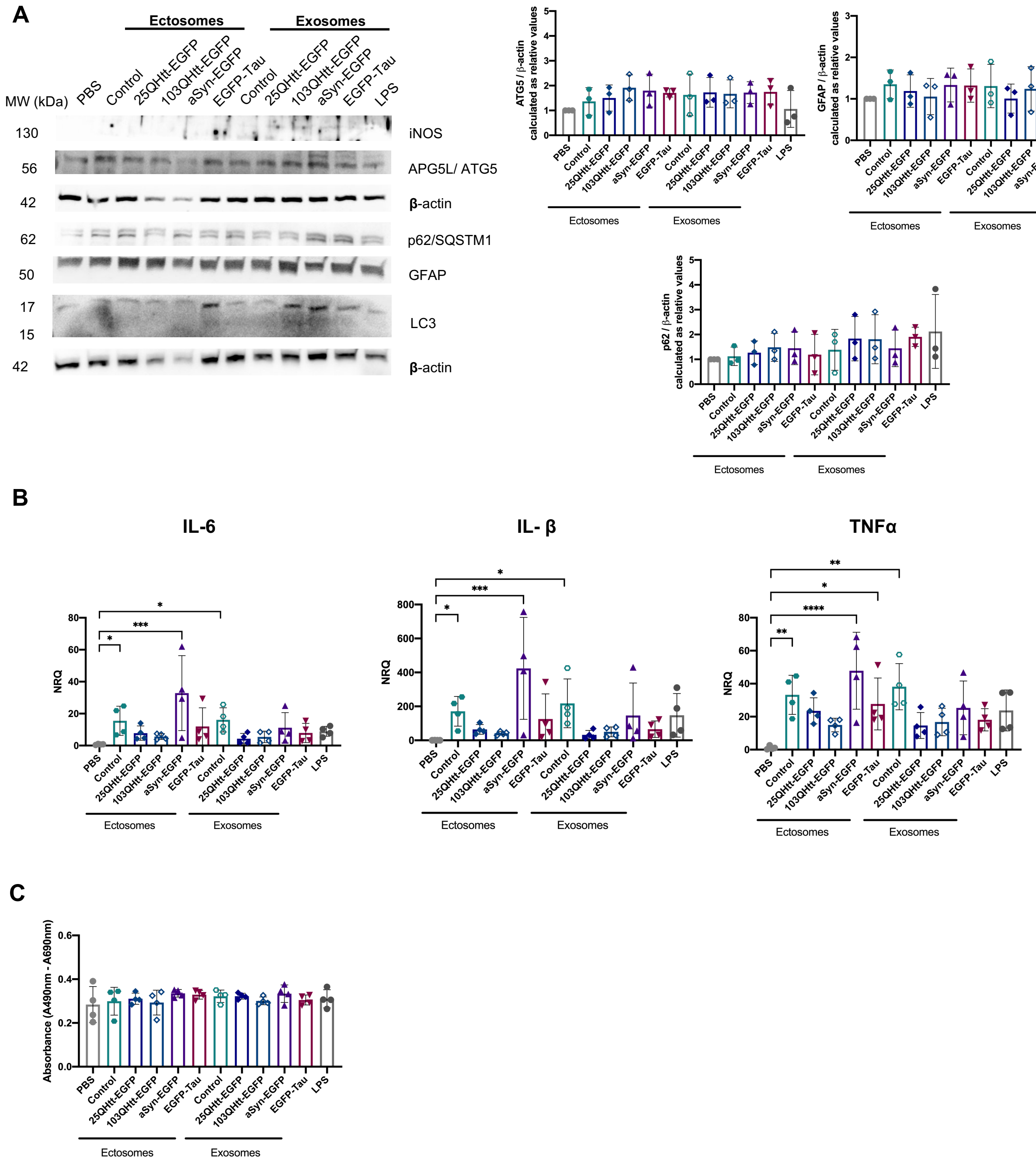


**Supplementary Figure 7. Ectosomes and exosomes containing disease-related proteins are internalized by astrocytic cells.**

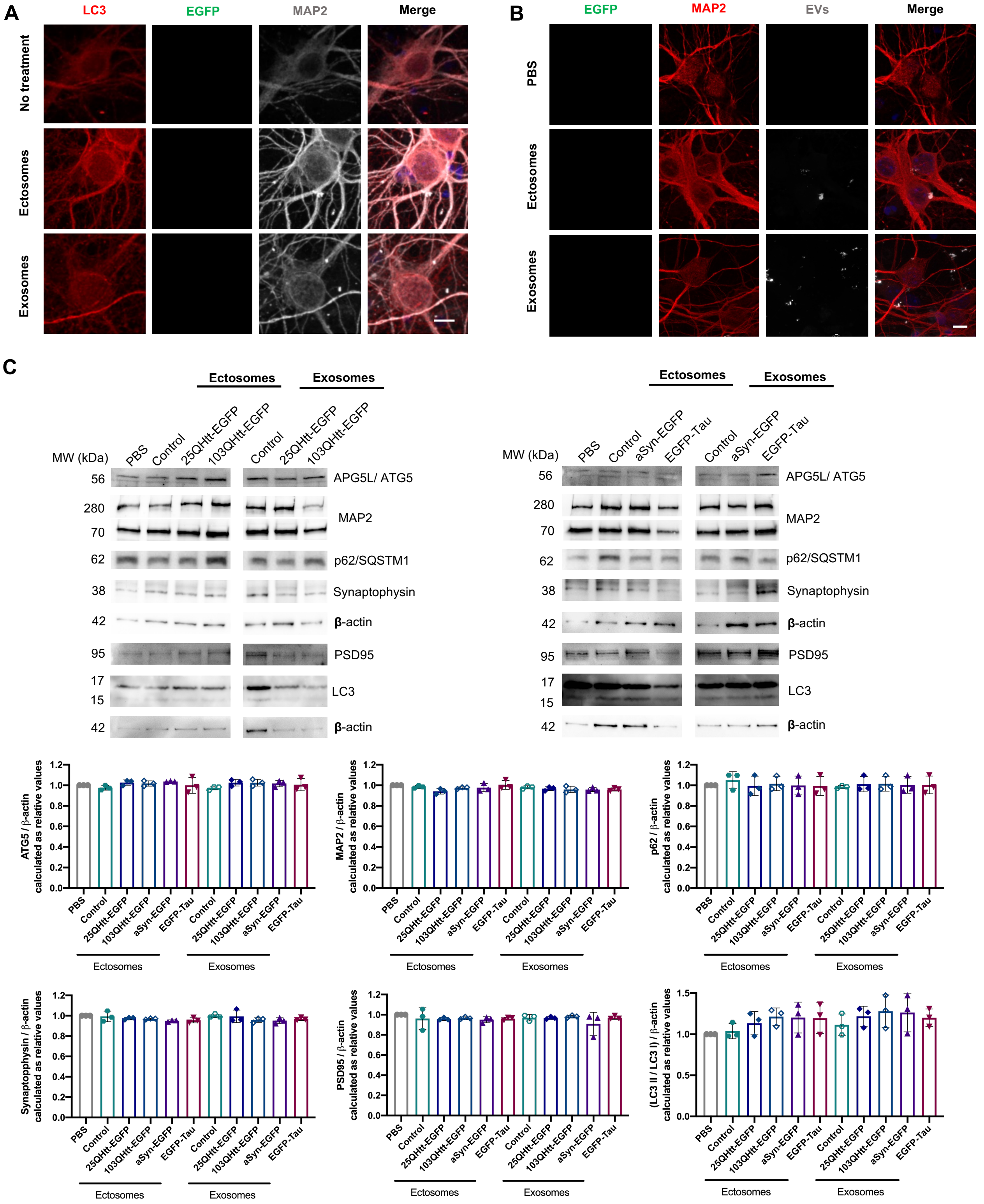


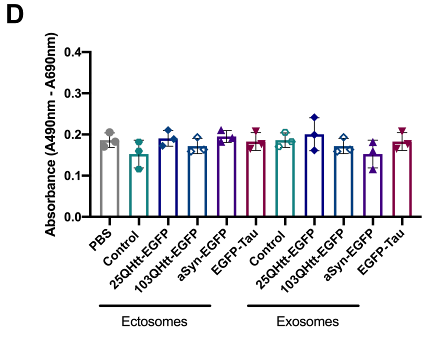


**Supplementary Figure 8. Ectosomes and exosomes containing disease-related proteins are internalized by cortical neurons.**
